# Tumor control of lysosomal acidification promotes lipoprotein assimilation and ferroptosis resistance

**DOI:** 10.64898/2026.08.25.747075

**Authors:** Renfei Wu, Sheng-Chieh Hsu, Lingjie Sang, Miaomiao Yu, Yoon Jung Kim, Mangyu Choe, Caroline Hauer, Ling Cai, Ariella B. Hanker, Isaac S. Chan, Hijai R. Shin, Javier Garcia-Bermudez

## Abstract

Lysosomes are acidic organelles that fuel cancer progression by facilitating nutrient acquisition and metabolic adaptation, yet the determinants through which cancer cells sustain specialized lysosomal functions are not fully delineated. Notably, assimilation of dietary antioxidants within lipoproteins, a lysosome-dependent process, protects tumors from ferroptosis, an oxidative form of cell death, raising the possibility that tumors evolve mechanisms to enhance this process. Here, we applied genetic screens to identify regulators of lysosome-dependent lipoprotein assimilation and ferroptosis resistance and identified ZNF217, a frequently amplified transcriptional regulator in human cancers, as a driver of tumor lysosomal function and ferroptosis resistance. ZNF217 promoted lipoprotein assimilation through transcriptional maintenance of RAB11FIP4, an endolysosomal protein. Loss of either ZNF217 or RAB11FIP4 impaired lysosomal acidification across multiple cancer types, leading to defective lipoprotein assimilation, increased lipid peroxidation, ferroptosis sensitivity, and impaired tumor growth. Mechanistically, RAB11FIP4 boosts lysosomal acidity through maintenance of RAB7A activity and proper assembly of the lysosomal V-ATPase complex. Finally, disruption of ZNF217 in breast cancer cell lines and patient-derived organoids, a tumor context linked to ZNF217 expression, reduced lysosomal acidity and impaired cancer growth through increased ferroptosis sensitivity. Together, we identify transcriptional regulation of lysosomal acidification as a key metabolic adaptation that enables extracellular antioxidant acquisition and tumor progression.

**SIGNIFICANCE:** Cancer cells exploit lysosomes for metabolic adaptation, but how tumors regulate lysosomal function to sustain nutrient assimilation remains poorly understood. We identify a ZNF217–RAB11FIP4 axis that maintains lysosomal acidification to promote lipoprotein-derived antioxidant assimilation and ferroptosis resistance, revealing transcriptional control of lysosomes as a tumor adaptation mechanism and pro-ferroptotic therapeutic vulnerability.

## INTRODUCTION

Lysosomes are central metabolic hubs^1^ that enable tumor adaptation to nutrient stress^2^. Beyond their classical degradative functions associated to nutrient scavenging^3,4^, lysosomes coordinate nutrient sensing^5^, intracellular trafficking^6^, and metabolic signaling pathways that support malignant growth. Tumors often exhibit increased lysosomal abundance and activity^7^, and disruption of lysosomal function impairs tumor progression across diverse cancer contexts^6,8,9^. Previous studies have largely focused on lysosomal biogenesis and autophagy as adaptive mechanisms in cancer, but less is known about how tumors regulate specific lysosomal functions that directly support metabolic fitness.

One lysosomal function that has emerged as particularly important for tumor metabolism is the assimilation of dietary lipids^10,11^. Many tumors increase uptake of circulating lipoproteins^12–15^, the carriers of lipids in circulation, a process that supports tumor aggressiveness. Following endocytosis, lipoprotein cargo is delivered to lysosomes, where acidic luminal pH enables cargo processing and release of metabolically useful lipids^16^. Lack of acidification impairs the effect of lysosomal lipases and inhibits dietary lipid utilization. Recent studies by our group and others have shown that lipoprotein-derived lipids, namely dietary vitamin E^15,17^, can function as a potent antioxidant that suppress lipid peroxidation and ferroptosis^18^, an important metabolic bottleneck of tumors^19^. Disruption of lipoprotein uptake impairs tumor growth by limiting acquisition of these lipid antioxidants and sensitizing tumors to ferroptosis^15^. Therefore, it is likely that maintenance of lysosomal acidity is necessary for tumors to efficiently assimilate dietary lipid antioxidants. However, the molecular pathways that sustain lysosomal acidification capacity compatible with lipoprotein assimilation in cancer cells remain unknown.

Here, using genetic screens designed to identify regulators of lysosome-dependent lipoprotein assimilation and ferroptosis resistance, we identify ZNF217 as a transcriptional regulator of lysosomal acidification in multiple cancer types. *ZNF217* is a frequently amplified locus across human cancers and has been implicated in tumor progression through transcriptional and epigenetic mechanisms^20,21^, but its role in lysosomal function has remained elusive. We further identify elevated expression of RAB11FIP4, an endolysosomal protein, as the primary ZNF217-regulated effector that drives cancer lysosomal function. Loss of RAB11FIP4 impairs the activity of the lysosomal GTPase RAB7A, which in turn lowers the lysosomal acidity milieu and disrupts proper assimilation of lipoproteins. Finally, we show that disruption of the ZNF217-RAB11FIP4 axis sensitizes tumors to ferroptosis and suppresses growth across multiple cancer types, including patient-derived organoids. Together, these findings establish transcriptional regulation of lysosomal acidification as a key metabolic adaptation that facilitates dietary antioxidant acquisition and enables tumor progression.

## RESULTS

### Genetic screens pinpoint *ZNF217* as necessary for lipoprotein assimilation and cancer ferroptosis resistance

A major functional advantage of lipoprotein uptake by tumors is the acquisition of dietary lipid antioxidants that protect them from lipid oxidative stress and ferroptosis. Efficient assimilation of lipoprotein-derived lipids requires two steps: endocytic uptake of lipoproteins, and delivery of cargo to functional, acidic lysosomes. Although the role of SREBPs as transcriptional regulators of lipoprotein uptake is well established^22^, less is known about the transcriptional networks involved in sustaining lysosomal acidity in tumors. Thus, we designed genetic screens to interrogate which transcriptional regulators are essential for lysosomal-dependent assimilation of lipid antioxidants and ferroptosis resistance in cancer cells.

We conducted two parallel CRISPR genetic screens using an sgRNA library targeting transcriptional and epigenetic regulators^23^ in cell lines with previously reported high lipoprotein uptake^15^ or enhanced lysosomal function^24^. In the first screen, Karpas299 lymphoma cells transduced with the sgRNA library were subjected to a proliferation-based screen in the presence or absence of sublethal concentrations of the ferroptosis inducing compound RSL3, a chemical inhibitor of the anti-ferroptotic protein GPX4^25^ (**Fig. 1A**). Surviving cells were collected after 14 population doublings, and sgRNA amplicons were sequenced to identify transcriptional and epigenetic regulators whose knockout sensitizes cancer cells to ferroptosis in vitro. This genetic screen identified Zinc Finger Protein 217 (*ZNF217*) as the top hit under RSL3 treatment, with knockout sensitizing cells to ferroptosis (**Fig. 1B; Supplementary Fig. 1A**). *RCOR1*, which encodes a transcriptional regulator reported to physically and functionally interact with ZNF217^26,27^, was the second strongest hit in the screen (**Fig. 1B; Supplementary Fig. 1A**). *SREBF1*, encoding one of the SREBP family members, also scored as essential, consistent with its role in lipid metabolism.

**Figure 1.**
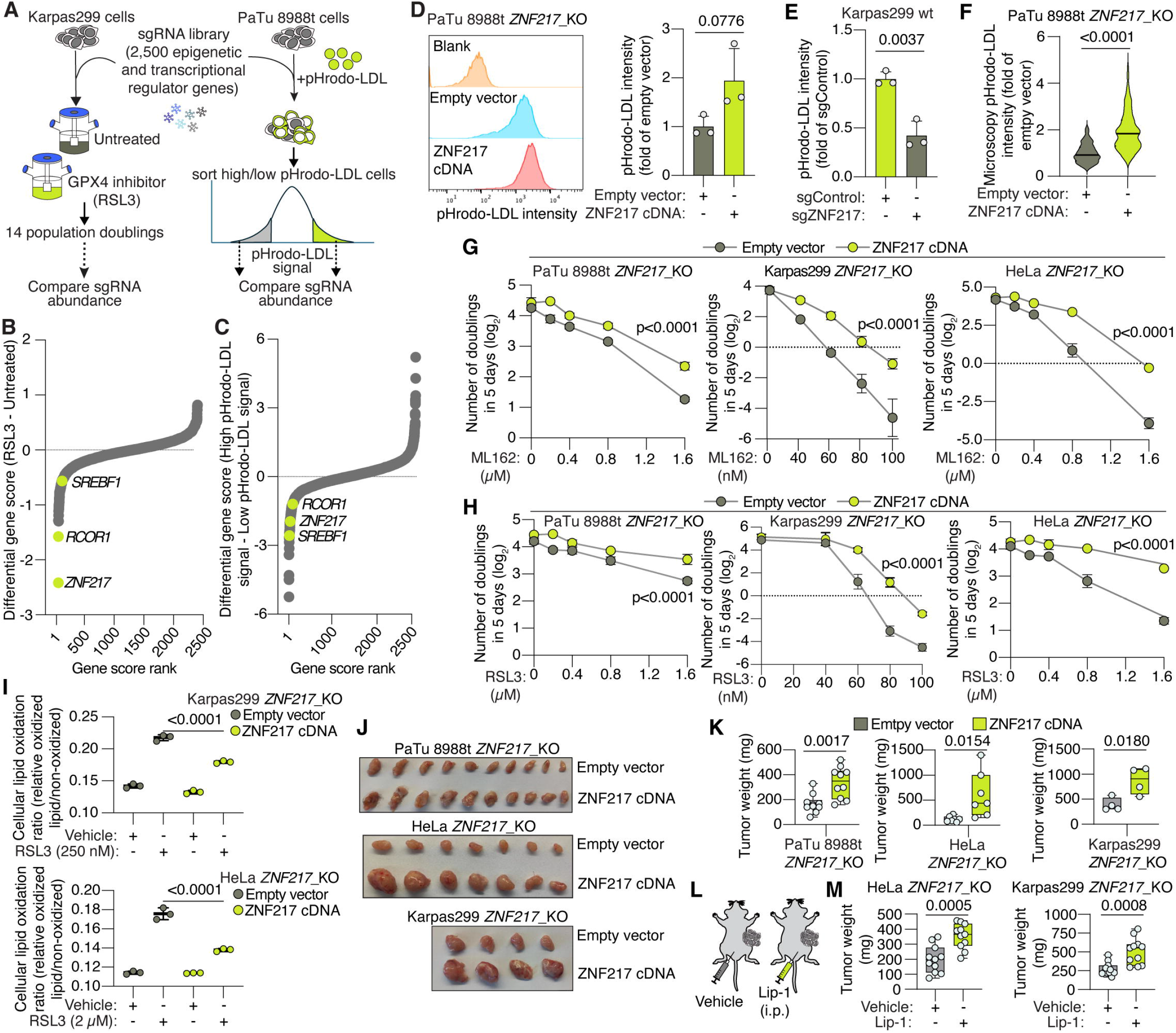
ZNF217 promotes lysosomal lipoprotein assimilation and ferroptosis resistance to sustain tumor growth. **A,** Schematic of focused CRISPR screens targeting 2,500 transcriptional and epigenetic regulators in Karpas299 cells cultured with or without RSL3 for 14 population doublings (left) or pHrodo-LDL-sorted PaTu 8988t cells (right). **B,** Differential gene scores from proliferation CRISPR screen in RSL3-treated versus untreated Karpas299 cells. **C,** Differential gene scores from pHrodo-LDL CRISPR screen comparing gene essentiality of PaTu 8988t cells with high pHrodo-LDL fluorescence to those with low pHrodo-LDL fluorescence. **D,** Representative flow cytometry histograms (left) and quantification (right) of pHrodo-LDL fluorescence in *ZNF217*_KO PaTu 8988t cells transduced with an empty vector or a ZNF217 cDNA. **E,** Quantification of pHrodo-LDL fluorescence in Karpas299 cells transduced with an sgControl or sgZNF217 in flow cytometry assays. **F,** Live-cell microscopy analysis of pHrodo-LDL fluorescence in the cells described in D. **G–H,** Number of doublings (log_2_) in 5 days of *ZNF217*_KO PaTu 8988t, Karpas299, and HeLa cells transduced with an empty vector or ZNF217 cDNA after treatment with the indicated concentrations of the GPX4 inhibitors ML162 (G) or RSL3 (H). **I,** Quantification of lipid oxidation in *ZNF217*_KO Karpas299 and HeLa cells transduced with an empty vector or ZNF217 cDNA after treatment with vehicle or the indicated concentrations of RSL3. **J–K,** Representative images (J) and tumor weights (K) of subcutaneous tumors formed by *ZNF217*_KO PaTu 8988t (n = 10), HeLa (n = 7), and Karpas299 (n = 4) cells transduced with an empty vector or ZNF217 cDNA. **L,** Schematic depicting the liproxstatin-1 (Lip-1) supplementation approach in tumor-bearing mice. **M,** Tumor weights from mice implanted with HeLa (n = 6) or Karpas299 (n = 6) *ZNF217*_KO cells under supplementation with vehicle or Lip-1. In vitro data (D–I) are mean ± s.d.; n = 3 biologically independent experiments. Tumor weights in K and M are shown as box plots; the center line indicates the mean, the box limits indicate the first and third quartiles, and whiskers indicate the range. Statistics were performed using two-sided unpaired t tests unless otherwise indicated. Data in G–I were analyzed by two-way ANOVA followed by Šídák’s multiple-comparison test.

In parallel, PaTu 8988t pancreatic ductal adenocarcinoma (PDAC) cancer cells, a cancer type characterized by enhanced lysosomal nutrient acquisition^3,24^, were incubated with LDL conjugated to pHrodo (pHrodo-LDL), an acidic pH-sensitive fluorogenic probe that fluoresces upon lysosomal delivery^28^. We first validated that the fluorescence readout was sensitive to lysosomal pH, as inhibition of the lysosomal V-ATPase with bafilomycin A1 (BafA1) decreased cellular pHrodo-LDL signal in flow cytometry assays (**Supplementary Fig. 1B**). PaTu 8988t cells transduced with the sgRNA library were subsequently sorted by flow cytometry (**Fig. 1A**). We isolated knockout cells with the highest (top 5% fluorescence) and lowest (bottom 5% fluorescence) pHrodo-LDL signal. sgRNA amplicons from each population were sequenced and sgRNA abundance compared to identify genes regulating delivery of LDL cargo to acidic lysosomes. This screen also identified *ZNF217*, *RCOR1*, and *SREBF1* among the genes whose loss decreased pHrodo-LDL signal (**Fig. 1C; Supplementary Fig. 1C**). Notably, the MiT/TFE family of transcription factors (*MITF*, *TFE3*, and *TFEB*), which control lysosomal biogenesis and function in cancer^24,29^, were not among the top hits, indicating that our genetic screen captures changes in lysosomal assimilation of lipoproteins and not broader changes in lysosomal biogenesis.

Building upon these genetic screen results, we focused on *ZNF217*, which encodes an epigenetic regulator that has not been implicated in lysosomal biology before. We used CRISPR to generate *ZNF217* knockout (*ZNF217*_KO) cells in four cell lines with elevated lipoprotein uptake: PDAC PaTu 8988t, lymphoma Karpas299, lung adenocarcinoma NCI-H838, and HeLa cells (**Supplementary Fig. 1D**), prior to assessing pHrodo-LDL assimilation and sensitivity to ferroptosis. Loss of *ZNF217* reduced pHrodo-LDL fluorescence across cell lines (PaTu 8988t: 47% decrease; Karpas299: 58% decrease; HeLa: 38% decrease; **Fig. 1D, 1E and Supplementary Fig. 1E**), as measured by flow cytometry. These results were confirmed using live-cell microscopy (**Fig. 1F**).

Before being delivered to lysosomes, LDL particles are taken up by endocytosis. Importantly, the observed effect of ZNF217 on pHrodo-LDL assays was not driven by impaired LDL uptake, as flow cytometric uptake assays using LDL particles labeled with DiI, a pH-insensitive fluorescent dye^30^ (DiI-LDL), showed no change or only a modest decrease in *ZNF217*_KO cells relative to isogenic controls (**Supplementary Fig. 1F**). Thus, ZNF217 is required for efficient delivery of LDL cargo to acidic lysosomes.

Similarly, consistent with the RSL3-treated CRISPR screen results, *ZNF217*_KO cells were more sensitive to ferroptosis induction than isogenic counterparts expressing ZNF217 when treated with the GPX4 inhibitors ML162 and RSL3 (**Fig. 1G and 1H; Supplementary Fig. 1G**), or with erastin, a cystine uptake inhibitor that depletes glutathione and thus constitutes an orthogonal approach to chemically induce ferroptosis (**Supplementary Fig. 1H**). Given the established role of lipoproteins in promoting ferroptosis resistance via delivery of vitamin E^15,17^, together with the observed effects of ZNF217 on lysosomal lipoprotein assimilation, we next sought to determine whether these phenotypes are mechanistically linked.

### ZNF217 protects tumors from ferroptosis

Assimilation of lipoprotein-derived antioxidants promotes cancer cell resistance to ferroptosis, and all previous ferroptosis induction assays were performed in media supplemented with fetal bovine serum (FBS), which is rich in lipoproteins. To determine whether the effects of ZNF217 on ferroptosis resistance are mediated by lipoproteins, we repeated these proliferation experiments in the presence of ferroptosis inducers using media supplemented with lipoprotein-depleted serum (LPDS). Removal of lipoproteins abolished the effects of *ZNF217* loss in these assays and triggered both isogenic cell lines to equally respond to pharmacological ferroptosis induction (**Supplementary Fig. 2A**), suggesting that lipoprotein presence is required for this phenotype.

We next measured lipid oxidation using the lipid probe BODIPY-C11 in isogenic pairs under conditions of chemical GPX4 inhibition and found that *ZNF217*_KO cells accumulated higher levels of peroxidized lipids than ZNF217-expressing counterparts (**Fig. 1I**). This enhanced lipid oxidation rate was fully reversed by treatment with the lipid antioxidant ferrostatin-1 (Fer-1, **Supplementary Fig. 2B**). Importantly, performing this assay in lipoprotein-depleted media abolished the difference in cellular lipid oxidation between isogenic cells (**Supplementary Fig. 2C**). Thus, ZNF217 promotes cancer cell ferroptosis resistance in a lipoprotein-dependent manner.

Cancer anti-ferroptotic mechanisms, including lipoprotein assimilation, promote tumor growth and aggressiveness^15,31–35^. To test whether ZNF217 is required for tumor growth, we implanted human PaTu 8988t, HeLa, Karpas299, and NCI-H838 isogenic pairs with or without *ZNF217* expression subcutaneously into the flanks of immunodeficient mice. Across all four models, *ZNF217*_KO tumors were smaller than their ZNF217-expressing counterparts (**Fig. 1J-K; Supplementary Fig. 2D**), with tumor burden reduced by 50% in PaTu 8988t, 80% in HeLa, 55% in Karpas299, and 60% in NCI-H838 xenografts.

Lastly, to define whether this reduction in tumor growth was driven by increased lipid peroxidation and ferroptosis, we repeated tumor growth experiments using the two fastest-growing *ZNF217*_KO cell lines and their isogenic controls in mice treated with either the lipophilic radical-trapping antioxidant liproxstatin-1 (Lip-1), which rescues ferroptosis in vivo, or vehicle control (**Fig. 1L**). Lip-1 treatment stimulated the growth of Karpas299 and HeLa *ZNF217*_KO tumors by 1.9- and 1.8-fold, respectively (**Fig. 1M; Supplementary Fig. 2E**). In contrast, Lip-1 did not significantly affect the growth of ZNF217-expressing tumors (**Supplementary Fig. 2F and 2G**). Together, these experiments show that ZNF217 is necessary for tumors to resist ferroptosis in vivo.

**Figure 2.**
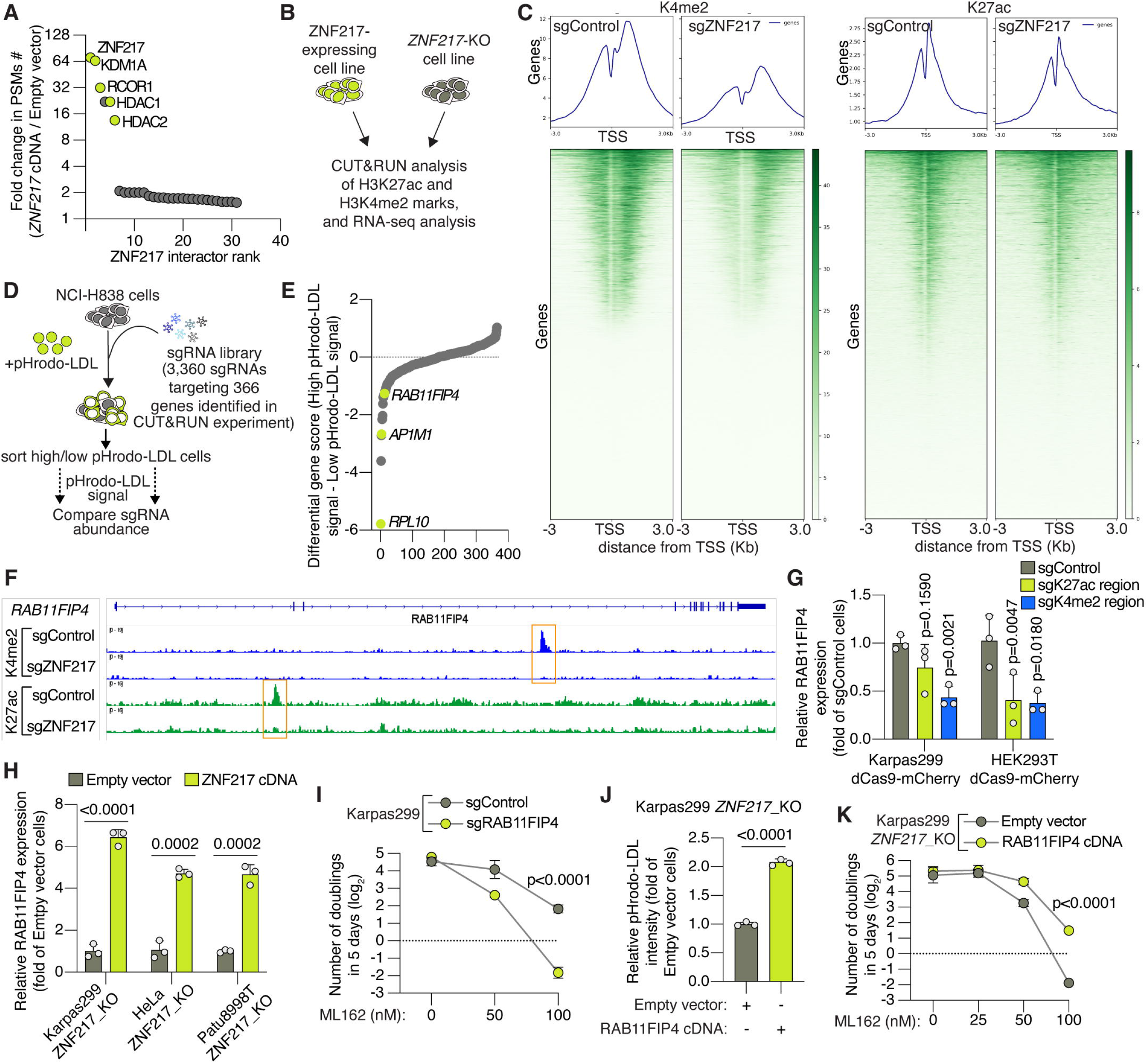
ZNF217 confers ferroptosis resistance by activating *RAB11FIP4* expression. **A,** Rank of the most enriched proteins identified in FLAG-ZNF217 pull-down assessed by proteomics in Karpas299 cells, based on the number of peptide-spectrum matches (PSMs) in the FLAG-ZNF217 group compared to the control group. A set of epigenetic regulating proteins are highlighted. **B,** Schematic of H3K4me2 and H3K27ac CUT&RUN analyses together with RNA-sequencing analysis in cancer cells with or without *ZNF217* expression. **C,** Global abundance of H3K4me2 and H3K27ac in Karpas299 cells transduced with sgZNF217 or a control sgRNA (sgControl). **D,** Schematic of a focused pHrodo-LDL CRISPR screen in NCI-H838 cells using 3,360 sgRNAs targeting 366 genes whose histone marks were found altered in CUT&RUN assays. **E,** Differential gene scores from pHrodo-LDL CRISPR screen comparing gene essentiality of NCI-H838 cells with high pHrodo-LDL fluorescence to those with low pHrodo-LDL fluorescence. **F,** H3K4me2 and H3K27ac CUT&RUN profiles at the *RAB11FIP4* locus in Karpas299 cells transduced with sgControl or sgZNF217. H3K4me2 and H3K27ac marks lost upon *ZNF217* loss are highlighted with orange rectangles. **G,** *RAB11FIP4* mRNA abundance in Karpas299 and HEK293T cells stably expressing dCas9 and transduced with control sgRNAs or sgRNAs targeting the H3K4me2- or H3K27ac-enriched regions shown in F. **H,** Relative *RAB11FIP4* mRNA expression in *ZNF217*_KO Karpas299, HeLa, and PaTu 8988t cells expressing an empty vector or ZNF217 cDNA. **I,** Number of doublings (log_2_) in 5 days of Karpas299 cells transduced with sgControl or sgRAB11FIP4 following treatment with the indicated concentrations of ML162. **J,** Quantification of pHrodo-LDL fluorescence in *ZNF217*_KO Karpas299 cells expressing empty vector or RAB11FIP4 cDNA. **K,** Number of doublings (log_2_) in 5 days of *ZNF217*_KO Karpas299 cells expressing empty vector or RAB11FIP4 cDNA following treatment with the indicated concentrations of ML162. Data in G–K are mean ± s.d.; n = 3 biologically independent experiments. Statistics were performed using two-sided unpaired t tests unless otherwise indicated. Data in I and K were analyzed by two-way ANOVA followed by Šídák’s multiple-comparison test. Panel F shows representative CUT&RUN tracks.

### ZNF217 promotes ferroptosis resistance in cancer cells through transcriptional upregulation of RAB11FIP4

ZNF217 is a protein that localizes to the nucleus (**Supplementary Fig. 3A**) previously reported to regulate gene expression through interactions with transcriptional regulators and epigenetic modifiers of histone marks^20,36^. To identify interacting partners of ZNF217 with potential functional relevance in our model, we expressed FLAG-tagged ZNF217 in Karpas299 cells and performed immunoprecipitation followed by mass spectrometry proteomics (IP-MS). This analysis identified multiple chromatin regulatory proteins among the top interacting partners (**Fig. 2A**), including RCOR1 (which also scored in our previous CRISPR screens; **Fig. 1B and Fig. 1C**), KDM1A, HDAC1, and HDAC2, raising the possibility that ZNF217 may function within epigenetic regulatory complexes in these cancer cells and consistent with previous reports.

To define how ZNF217 influences transcription-associated chromatin marks, we performed CUT&RUN, a chromatin profiling approach for mapping histone modifications, and quantified the abundance of two major histone marks, H3K4me2 and H3K27ac, in Karpas299 cells transduced with sgZNF217 or a control sgRNA (**Fig. 2B**). Global levels of both H3K4me2 and H3K27ac were reduced in sgZNF217-transduced cells (**Fig. 2C**). To generate an unbiased gene list for downstream screening, we performed differential peak analysis of H3K4me2 and H3K27ac, identifying 366 unique genes showing significant loss of either mark, or both (*Supplementary Table 1*). In parallel, we performed RNA-seq in NCI-H838 and Karpas299 isogenic cell line pairs with or without ZNF217 expression to generate a list of differentially expressed genes between genotypes. Notably, most transcriptional changes associated with *ZNF217* loss corresponded to decreased gene expression (*Supplementary Table 2*), consistent with the observed reduction in H3K4me2 and H3K27ac abundance, which are generally associated with transcriptional activation.

Because ZNF217 influences epigenetic histone marks across hundreds of loci, we next devised a CRISPR screening strategy to functionally prioritize candidate genes involved in lysosomal lipoprotein assimilation. We reasoned that a loss-of-function screen in parental ZNF217-expressing cells could identify genes whose disruption phenocopies the effects of *ZNF217* loss on pHrodo-LDL assimilation. We therefore constructed an sgRNA library targeting the 366 genes identified by CUT&RUN as having histone marks influenced by *ZNF217* expression, the majority of which were not included in the sgRNA library we used in the initial CRISPR screen, and performed a flow cytometry-based pHrodo-LDL screen in NCI-H838 cells (**Fig. 2D**). Analysis of gene essentiality in cell populations with high or low pHrodo-LDL fluorescence identified genes whose knockout reduced delivery of LDL cargo to acidic lysosomes (**Fig. 2E; Supplementary Fig. 3B**). Among these candidates, *AP1M1* and *RAB11FIP4* stood out based on their established roles in endocytosis^37^ and lysosomal biology^38^, respectively. Because the remaining hits largely converged on core gene expression and protein homeostasis pathways, such as ribosome biogenesis (*RPL10*), which are likely to influence multiple cellular processes beyond the endolysosomal system, we focused subsequent studies on AP1M1 and RAB11FIP4.

We then checked the expression of these three genes in our RNAseq datasets, and found that only *RAB11FIP4* mRNA levels were dependent on *ZNF217* expression (**Supplementary Fig. 3C**). These results suggest that changes in histone marks at the *RPL10* and *AP1M1* loci (**Supplementary Fig. 3D**) do not influence their transcription. In contrast, CUT&RUN analysis identified two major genomic regions within the *RAB11FIP4* locus that were marked by H3K4me2 or H3K27ac in ZNF217-expressing cells but not in *ZNF217*_KO cells (**Fig. 2F**). Furthermore, blocking modification of these regions by targeting a dead Cas9 to them via a specific sgRNA^39^ (**Supplementary Fig. 3E**) reduced *RAB11FIP4* mRNA abundance in two parental ZNF217-expressing cell lines (**Fig. 2G**), with the K4me2 mark having a stronger effect on *RAB11FIP4* expression. Consistent with these results, expression of a sgRNA-resistant ZNF217 cDNA in three KO cell lines recovered *RAB11FIP4* mRNA expression (**Fig. 2H**). Thus, spatially enriched histone marks in the *RAB11FIP4* gene are associated with increased expression, and are lost upon *ZNF217* loss.

Based on these observations, we focused on RAB11FIP4 for functional studies. First, we tested whether ZNF217 promotes LDL assimilation and ferroptosis resistance primarily through increasing *RAB11FIP4* expression. Knocking out *RAB11FIP4* in multiple cell lines (**Supplementary Fig. 4A and 4B**) sensitized ZNF217-expressing cells to chemical ferroptosis induction (**Fig. 2I; Supplementary Fig. 4C**), reduced pHrodo-LDL fluorescence (**Supplementary Fig. 4D**), and resulted on elevated cellular lipid peroxidation following GPX4 inhibition measured with the lipid peroxidation probe LiperFluo (**Supplementary Fig. 4E**). Moreover, ectopic expression of RAB11FIP4 cDNA under a constitutively active promoter in *ZNF217*_KO cells increased pHrodo-LDL fluorescence (**Fig. 2J**) and enhanced resistance to two different GPX4 inhibitors (**Fig. 2K; Supplementary Fig. 4F**), thereby phenocopying the effects of ZNF217 expression. Similar to loss of *ZNF217*, RAB11FIP4 expression did not significantly alter DiI-LDL uptake (**Supplementary Fig. 4G and 4H**). These experiments suggest that maintenance of *RAB11FIP4* expression is the primary driver of ZNF217-dependent lipoprotein-mediated ferroptosis resistance.

**Figure 3.**
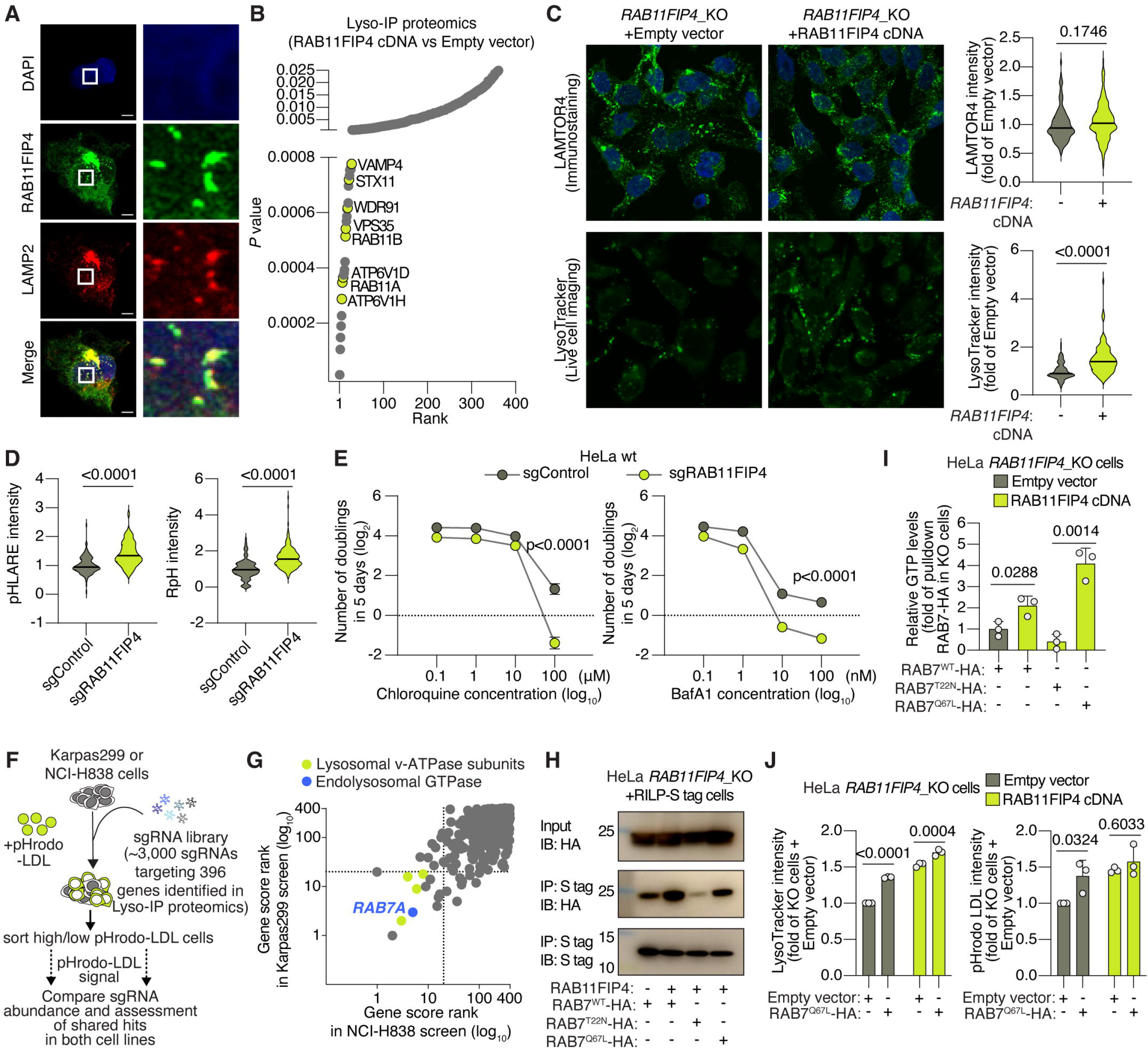
RAB11FIP4 promotes cancer lysosomal acidification. **A,** Representative immunofluorescence images showing colocalization of RAB11FIP4 and LAMP2. Scale bar, 5 μm. **B,** Differentially enriched proteins in lysosomal fractions of Karpas299 *RAB11FIP4*_KO cells expressing RAB11FIP4 cDNA compared to lysosomes of empty vector-expressing cells identified by Lyso-IP proteomics. Proteins associated with lysosomal acidification or maturation are highlighted. **C,** Representative images (left) and quantification (right) of LAMTOR4 immunostaining in fixed cells (top), and LysoTracker fluorescence assessed by live-cell imaging (bottom) in *RAB11FIP4*_KO PaTu 8988t cells expressing an empty vector or RAB11FIP4 cDNA. **D,** Quantification of lysosomal pH using genetically encoded pHLARE and RpH probes and live-cell imaging in HeLa cells transduced with sgControl or sgRAB11FIP4. **E,** Number of doublings (log_2_) in 5 days of HeLa cells transduced with sgControl or sgRAB11FIP4 following treatment with the indicated concentrations of chloroquine (CQ, left) or bafilomycin A1 (BafA1, right). **F,** Schematic of focused fluorescence-based pHrodo-LDL CRISPR screens in Karpas299 and NCI-H838 cells using approximately 3,000 sgRNAs targeting 396 genes identified by Lyso-IP proteomics. **G,** Quadrant map of overlapping genes from the two genetic screens in F. Shared hits include lysosomal V-ATPase subunit genes (green) and the small GTPase *RAB7A* (blue). **H,** Immunoblot of RAB7-HA following S-tag pulldown in HeLa *RAB11FIP4*_KO or *RAB11FIP4*-expressing cells expressing the indicated HA-tagged RAB7 cDNAs and transiently expressing RILP-S-tag. **I,** Quantification of GTP bound to indicated RAB7-HA protein isoforms following anti-HA immunoprecipitation from HeLa *RAB11FIP4*_KO expressing an empty vector or RAB11FIP4 cDNA. **J,** Flow cytometry analysis of LysoTracker (left) and pHrodo-LDL (right) fluorescence in HeLa *RAB11FIP4*_KO cells transduced with an empty vector or RAB11FIP4 cDNA and transiently expressing constitutively GTP-bound RAB7^Q67L^. Data in C–E and I–J are mean ± s.d.; n = 3 biologically independent experiments. Panel A and H are representative of three biologically independent experiments. Statistics were performed using two-sided unpaired t tests unless otherwise indicated. Data in E and J were analyzed by two-way ANOVA followed by Šídák’s multiple-comparison test. Panel B summarizes results from three Lyso-IP proteomic experiments.

**Figure 4.**
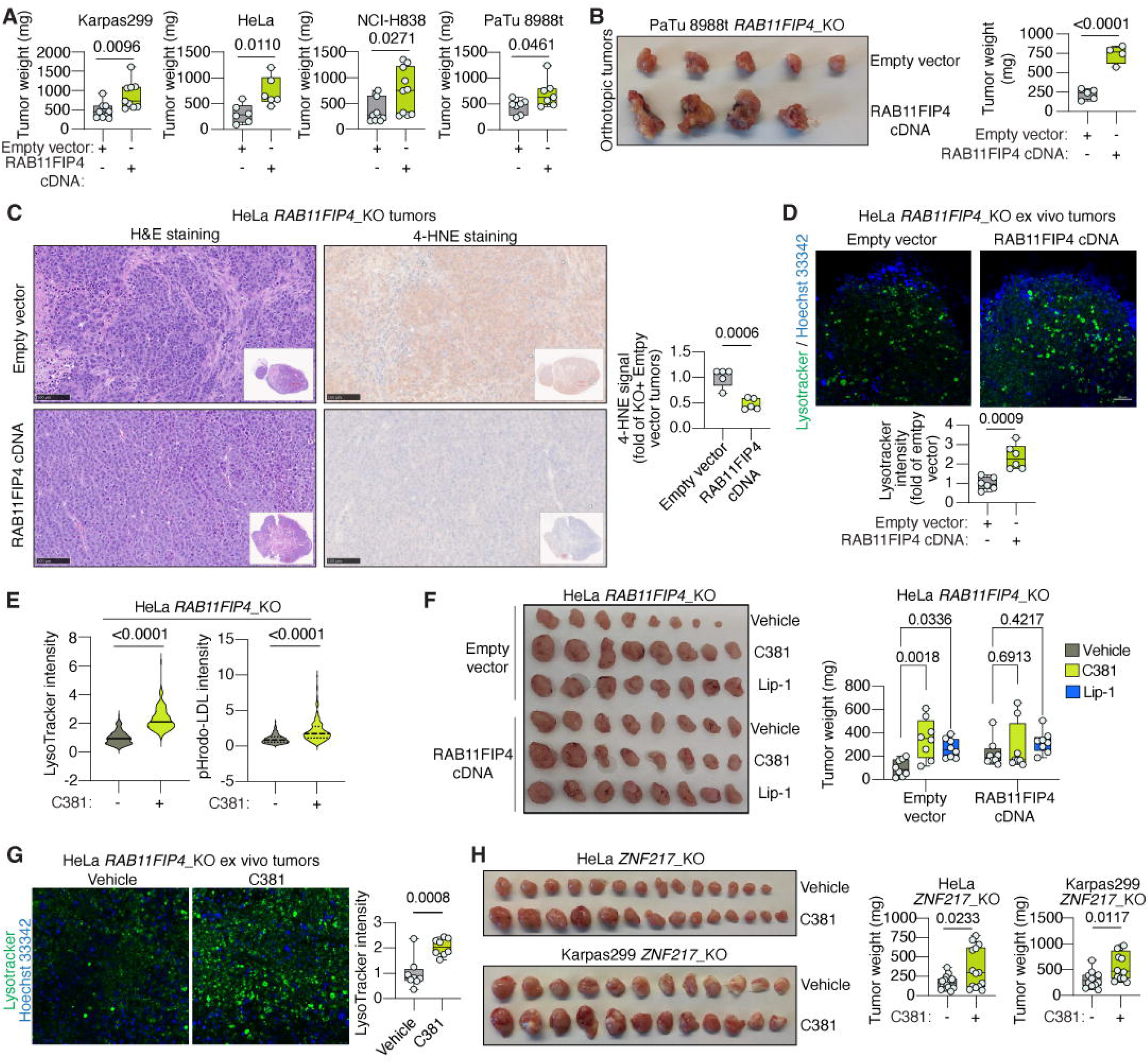
Lysosomal acidity maintenance by the ZNF217-RAB11FIP4 axis promotes tumor aggressiveness. **A,** Tumor weights of subcutaneous tumors formed by Karpas299 (n = 9), HeLa (n = 6), NCI-H838 (n = 10), and PaTu 8988t (n = 7) *RAB11FIP4*_KO cells expressing an empty vector or RAB11FIP4 cDNA. **B,** Representative image (left) and tumor weights (right) of orthotopic tumors formed by pancreatic cancer PaTu 8988t *RAB11FIP4*_KO cells expressing an empty vector (n = 5) or RAB11FIP4 cDNA (n = 4). **C,** Representative hematoxylin and eosin (H&E) staining and 4-hydroxy-nonenal (4-HNE) immunohistochemistry (left) and signal quantification (right) of tumors arising from HeLa *RAB11FIP4*_KO cells expressing an empty vector or RAB11FIP4 cDNA (n = 5 tumors per group). **D,** Representative images (top) and quantification (bottom) of LysoTracker staining in ex vivo tumor slices arising from HeLa *RAB11FIP4*_KO expressing an empty vector or RAB11FIP4 cDNA (n = 6 tumors per group). Hoechst 33342 was used as a nuclear counterstain. **E,** Quantification of LysoTracker (left) and pHrodo-LDL (right) fluorescence assessed by live-cell imaging in HeLa *RAB11FIP4*_KO cells following treatment with vehicle or C381 (10 μM) for 24 hours. **F,** Representative images (left) and tumor weights (right) of subcutaneous tumors formed by HeLa *RAB11FIP4*_KO cells expressing an empty vector or RAB11FIP4 cDNA after treatment with vehicle, C381 (30 mg kg⁻¹, twice weekly), or Lip-1 (10 mg kg⁻¹, every two days) for 3 weeks (n = 8 tumors per treatment group). **G,** Representative LysoTracker staining (left) and quantification (right) of ex vivo tumor slices from vehicle- or C381-treated HeLa *RAB11FIP4*_KO tumors (n = 8 tumors per group). **H,** Representative images (left) and tumor weights (right) of subcutaneous HeLa (n = 7) or Karpas299 (n = 6) *ZNF217*_KO tumors following treatment with vehicle or C381 (30 mg kg⁻¹, twice weekly) for 3 weeks. Tumor weights in A, B, D, F, and H are shown as box plots; the center line indicates the mean, the box limits indicate the first and third quartiles, and whiskers indicate the range. Panels C and G show representative images only. Statistics for panel F were performed using two-way ANOVA followed by Šídák’s multiple-comparison test. All other comparisons were performed using two-sided unpaired t tests unless otherwise indicated.

### RAB11FIP4 is necessary for maintenance of lysosomal acidity

Although RAB11FIP4 has previously been linked to lysosomal function^38^, its specific roles in lysosomal biology and efficient lipoprotein assimilation are poorly understood. Immunostaining in parental HeLa cells revealed that RAB11FIP4 colocalizes with LAMP2, an endolysosomal marker, suggesting RAB11FIP4 localizes to the endolysosomal compartment (**Fig. 3A**).

To unbiasedly define how loss of *RAB11FIP4* influences lysosomal composition, we performed rapid lysosome immunopurification^40^ (Lyso-IP, **Supplementary Fig. 5A**) followed by quantitative proteomics (*Supplementary Table 3*). Multiple proteins were significantly depleted in lysosomes isolated from *RAB11FIP4*_KO cells (**Fig. 3B; Supplementary Fig. 5B**), including endolysosomal RAB GTPases, and components of the SNARE fusion machinery^41^. Notably, multiple V1 subunits of the lysosomal V-ATPase, which generates the proton gradient required for acidic lysosomal pH, were consistently reduced at the protein level in KO cells compared to control cells (ATP6V1A, ATP6V1B2, ATP6V1G1, and ATP6V1H; **Supplementary Fig. 5B**).

The reduced abundance of multiple V1 subunits in Lyso-IP proteomics prompted us to test whether assembly of the lysosomal V-ATPase complex^42,43^ was impaired in *RAB11FIP4*_KO cells. We immunoprecipitated ATP6V0A1, a V0 subunit, and assessed co-immunoprecipitation of ATP6V1A, a V1 subunit, as a proxy for V0-V1 assembly. Loss of *RAB11FIP4* reduced association between V0 and V1 subunits, suggesting impaired assembly of the lysosomal V-ATPase complex (**Supplementary Fig. 5C**).

Proper V0-V1 assembly is instrumental for the ability of the V-ATPase to maintain lysosomal acidity. We next directly assessed lysosomal pH in cells with or without *RAB11FIP4* expression using three orthogonal approaches: LysoTracker, a lysosome-targeted pH-sensitive fluorogenic dye; pHLARE, a genetically encoded lysosomal pH reporter^44^; and RpH, another genetically encoded reporter which directly measures lysosomal luminal pH^45^. *RAB11FIP4*_KO cells exhibited reduced lysosomal acidification compared with RAB11FIP4-expressing cells in LysoTracker assays using two orthogonal live-cell imaging approaches (**Fig. 3C; Supplementary Fig. 6A and 6B**), and in live-cell imaging experiments using both genetically encoded lysosomal pH reporters (**Fig. 3D**). Importantly, these changes in lysosomal pH occurred without significant differences in lysosome abundance, as assessed by LAMTOR4 immunostaining (**Fig. 3C**). Moreover, they were specific to the endolysosomal compartment, as measurement of cytosolic pH using the probe BCECF revealed no changes between cells with or without *RAB11FIP4* expression (**Supplementary Fig. 6C**). Finally, loss of *ZNF217* phenocopied these defects in lysosomal acidification (**Supplementary Fig. 6D and 6E**), consistent with the loss in *RAB11FIP4* expression previously observed in this genetic context.

To more precisely quantify the magnitude of this defect, we generated lysosomal pH calibration curves using LysoSensor^46^ in parental cells, *RAB11FIP4*_KO cells, and KO cells transduced with a sgRNA resistant RAB11FIP4 cDNA. While LysoTracker simply reports whether lysosomes are acidic, LysoSensor is a probe that allows for precise intra-lysosomal pH quantification. As a reference for complete inhibition of lysosomal acidification, we included treatment with BafA1, which fully inhibits V-ATPase activity. *RAB11FIP4*_KO cells exhibited an approximately 0.4-unit increase in lysosomal pH relative to isogenic cells expressing RAB11FIP4 (**Supplementary Fig. 6F**), whereas BafA1 treatment increased lysosomal pH to ∼8.0. Because the pH scale is of logarithmic nature, this 0.4-unit increase corresponds to an approximately 2.5-fold reduction in lysosomal proton concentration, suggesting a partial rather than complete defect in lysosomal acidification.

Consistent with this, mTORC1 signaling, which is tightly linked to lysosomal acidity^47^, was decreased but not abolished in *RAB11FIP4*_KO cells. Phospho-S6K levels, which can be used as a proxy of mTORC1 signaling, were reduced by approximately 50% under refeeding conditions while remaining readily detectable (**Supplementary Fig. 6G and 6H**). Finally, to determine whether these pH changes have functional consequences for cell fitness, we inhibited lysosomal acidification using BafA1 or chloroquine^48^ and assessed proliferation (**Fig. 3E; Supplementary Fig. 6I**). *RAB11FIP4*_KO cells were more sensitive to both compounds despite proliferating similarly to control cells under vehicle treatment. Therefore, the partial loss of lysosomal acidification caused by *RAB11FIP4* loss is sufficient to sensitize to pharmacological lysosomal inhibition and to impair lipoprotein assimilation and resistance to ferroptosis, but is compatible with unaltered baseline proliferation under nutrient-rich culture conditions.

Although RAB11FIP4 has been implicated in lysosomal storage disease^38^, it has not been previously linked to lysosomal acidity maintenance and the mechanism by which it supports lysosomal function is unknown. Small GTPases are central regulators of endolysosomal trafficking, dictating vesicular transport and lysosomal maturation required for lysosomal function and acidification^49^. RAB11FIP4 lacks intrinsic GTPase activity and has been proposed to function through interactions with RAB11^50^, a GTPase involved in vesicular trafficking. RAB11 protein levels were higher in lysosomes purified from RAB11FIP4-expressing cells than in those from KO counterparts (**Supplementary Fig. 7A**). However, whether RAB11FIP4 sustains lysosomal acidification through RAB11, other endolysosomal GTPases, or a distinct, GTPase-independent functional node is unclear.

To unbiasedly identify lysosomal functional modules that regulate acidity and lipoprotein assimilation downstream of RAB11FIP4, we constructed an sgRNA library targeting the ∼400 proteins identified in our Lyso-IP proteomics dataset, reasoning that relevant RAB11FIP4-associated factors would be represented. We transduced NCI-H838 and Karpas299 cells with this library and performed flow cytometry-based CRISPR screens using pHrodo-LDL signal as a functional readout (**Fig. 3F**), similar to the previous genetic screens that identified *ZNF217* and *RAB11FIP4*. After defining gene essentiality scores (**Supplementary Fig. 7B and 7C**), we prioritized genes whose loss reduced pHrodo-LDL signal in both cell lines (**Fig. 3G**). Among the strongest shared hits were four V1 subunits of the lysosomal V-ATPase (*ATP6V1A*, *ATP6V1B2*, *ATP6V1G1*, and *ATP6V1H*), which were also depleted in lysosomes from *RAB11FIP4*_KO cells in the previous proteomics experiment (**Supplementary Fig. 5B**) and served as a positive control for the screen given their essential role in promoting lysosomal acidity. In addition to these V-ATPase components, *RAB7A* emerged as a particularly compelling phenotype-driving candidate because of its established roles in lysosomal biology and positioning^51^.

We next tested whether loss of *RAB11FIP4* results on impaired RAB7A activity. RAB7A cycles between an inactive GDP-bound state and an active GTP-bound membrane-associated state that recruits downstream effectors such as Rab-Interacting Lysosomal Protein^52^ (RILP), to drive late endosomal and lysosomal trafficking and maturation. Fortunately, inactivating and activating mutations of RAB7A have been previously identified^53^. We ectopically expressed wild-type RAB7A together with constitutively active (Q67L – stays in the GTP-bound state) or dominant-negative (T22N) mutants in cells with or without RAB11FIP4 expression (**Fig. 3H**) and assessed activation state using RILP binding and GTP loading as readouts. Co-immunoprecipitation experiments revealed reduced association between RAB7A and RILP in *RAB11FIP4*_KO cells relative to controls (**Fig. 3H**), suggesting lower GTP-bound state in KO cells compared to RAB11FIP4-expressing ones. We confirmed that RAB7A has reduced GTP loading when RAB11FIP4 is lost by pulling down ectopically expressed wild-type RAB7A and measuring GTP levels. This assay revealed 52% lower GTP loading in *RAB11FIP4*_KO cells (**Fig. 3I**). Expression of the dominant-negative T22N mutant reduced RAB7A-RILP association and GTP loading, whereas the constitutively active Q67L mutant maintained stable interaction with RILP and elevated GTP-bound levels (**Fig. 3H and 3I**; **Supplementary Fig. 7D**), confirming the mutant phenotypes and the robustness of the approach. Therefore, RAB11FIP4 expression promotes the GTP-bound, active state of RAB7A.

Lastly, we tested whether GTP-bound, active RAB7A is necessary and sufficient for maintenance of lysosomal acidity and lipoprotein uptake in cancer cells. Because RAB7A is broadly required for lysosomal function and its complete loss is poorly tolerated in cells, we instead silenced CCZ1^54^, a component of the guanine nucleotide exchange machinery that activates RAB7A by promoting GDP-to-GTP exchange. *CCZ1* silencing (**Supplementary Fig. 7E**) reduced both LysoTracker intensity and pHrodo-LDL signal in RAB11FIP4-expressing cells (**Supplementary Fig. 7F**), thus phenocopying loss of *RAB11FIP4*.

Conversely, expression of constitutively GTP-bound, active RAB7A (Q67L) increased both LysoTracker and pHrodo-LDL signals in *RAB11FIP4*_KO cells to levels similar to in RAB11FIP4-expressing cells, while having minimal effects in RAB11FIP4-expressing controls (**Fig. 3J**), suggesting that RAB7A activation is sufficient to rescue the loss of lysosomal acidity triggered by *RAB11FIP4* loss. Together, these experiments show that RAB7A activity is necessary and sufficient to maintain lysosomal-dependent lipoprotein assimilation, and that loss of *RAB11FIP4* expression is linked to defective RAB7A activation and reduced lysosomal acidity.

### Lysosomal acidity maintenance by the ZNF217-RAB11FIP4 axis is necessary for tumor growth

In previous experiments, we found that loss of *ZNF217* impairs tumor growth and that chemical inhibition of ferroptosis with Lip-1 rescues these growth defects. We next defined whether RAB11FIP4 similarly regulates tumor growth. Isogenic pairs with or without *RAB11FIP4* expression from the same four cell lines used for ZNF217 studies (Karpas299, HeLa, NCI-H838, and PaTu 8988t) were implanted subcutaneously into mice and tumor weights assessed at endpoint. Loss of *RAB11FIP4* reduced tumor growth across all four models (**Fig. 4A; Supplementary Fig. 8A**). This phenotype was not restricted to subcutaneous models, as orthotopic implantation of PDAC PaTu 8988t *RAB11FIP4*_KO tumors resulted in a 70% reduction in pancreatic tumor burden compared with controls (**Fig. 4B**).

This impaired tumor growth upon *RAB11FIP4* loss correlated with increased lipid peroxidation and ferroptosis in vivo. We assessed tissue lipid oxidation via immunohistochemistry of 4-hydroxy-nonenal^55^ (4-HNE), a by-product of lipid peroxidation reactions and marker of ferroptosis in vivo. HeLa *RAB11FIP4*_KO tumors had enhanced levels of 4-HNE relative to isogenic controls (**Fig. 4C**). Consistent with our prior in vitro observations, these *RAB11FIP4*_KO tumor cells also exhibited a 57% reduction in LysoTracker intensity when analyzed ex vivo (**Fig. 4D**), supporting impaired tumor lysosomal acidification. Thus, loss of *RAB11FIP4* in tumors recapitulates the in vitro observations of increased lipid peroxidation and decreased lysosomal acidity.

We next attempted to pharmacologically rescue both lysosomal acidity and ferroptosis in tumors growing in mice to test the contribution of both phenotypes in cells with or without *RAB11FIP4* expression. For lysosomal acidity, we used C381, a lysosome-targeted small molecule that promotes lysosomal acidification^56^ and has previously been used to enhance clearance of toxic protein aggregates^57^. Treatment of *RAB11FIP4*_KO cells with C381 in vitro increased both LysoTracker and pHrodo-LDL signals (**Fig. 4E; Supplementary Fig. 8B and 8C**), validating the previously reported restoration of lysosomal function and its effect on lipoprotein assimilation, respectively.

Next, we repeated tumor growth experiments in mice receiving intraperitoneal injections of either Lip-1, C381, or vehicle control (**Supplementary Fig. 8D**). Supplementation of tumor-bearing mice with the lipid antioxidant Lip-1 selectively stimulated growth of *RAB11FIP4*_KO tumors while having minimal effects on RAB11FIP4-expressing counterparts (**Fig. 4F; Supplementary Fig. 8E**). C381 treatment rescued the growth defect of HeLa and PaTu 8988t *RAB11FIP4*_KO tumors (**Fig. 4F; Supplementary Fig. 8F**). Importantly, C381 supplementation reduced 4-HNE accumulation in *RAB11FIP4*_KO tumors (**Supplementary Fig. 8G**), suggesting that restoration of lysosomal acidity decreases lipid peroxidation in vivo. Moreover, C381 restored LysoTracker intensity in tumor cells analyzed ex vivo (**Fig. 4G**), confirming its delivery and on-target effect on lysosomal acidity in these tumors.

Lastly, because loss of *ZNF217* decreases *RAB11FIP4* levels and lysosomal acidity, we also tested whether pharmacologically increasing lysosomal acidity in *ZNF217*_KO tumors similarly rescued growth. Indeed, C381 treatment enhanced growth of *ZNF217*_KO tumors compared with vehicle-treated controls (**Fig. 4H**). In summary, these experiments identify impaired lysosomal acidification as a primary driver of the reduced tumor aggressiveness triggered by disruption of the ZNF217-RAB11FIP4 axis.

### ZNF217 promotes growth of breast cancer tumors and patient-derived organoids

Because ZNF217 plays a well-established role in breast cancer pathogenesis^20^, we chose this disease model to investigate its pre-clinical link to lysosomal function. In line with prior studies^58,59^, *ZNF217* copy-number gain directly tracked with increased gene expression in breast tumors and cell lines (**Fig. 5A**). While ZNF217 overexpression is a recognized driver of breast cancer progression and treatment resistance^20^, its role in lysosomal biology has not been examined in this tumor type. Therefore, we evaluated the impact of ZNF217 on tumor aggressiveness and lysosomal acidity using breast cancer cell lines and patient-derived organoids (PDOs).

**Figure 5.**
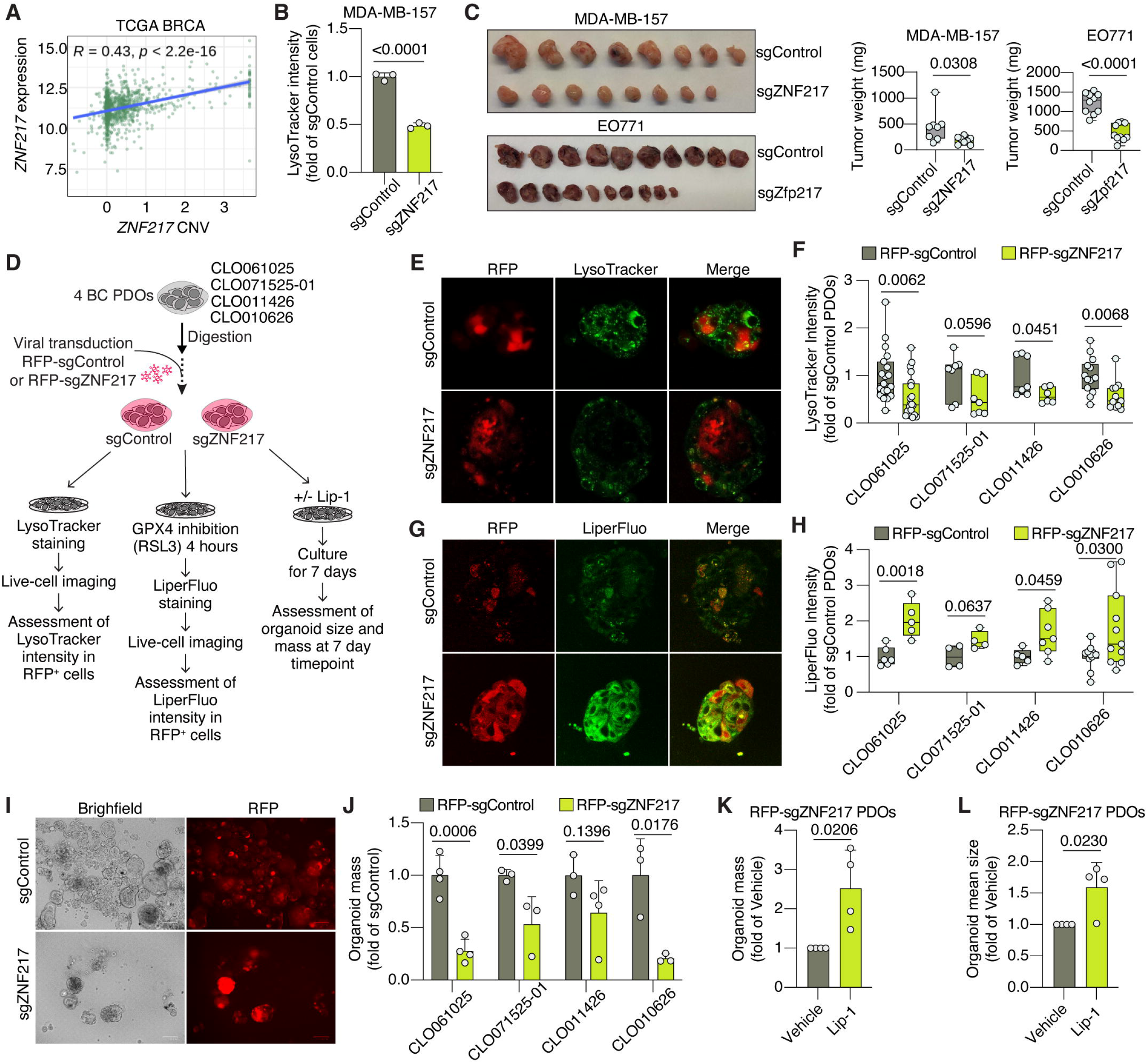
ZNF217 promotes lysosomal activity and growth of breast tumors and patient-derived organoids. **A,** Correlation between *ZNF217* copy number and *ZNF217* expression in the TCGA BRCA dataset. **B,** Flow cytometry analysis of LysoTracker fluorescence in MDA-MB-157 cells transduced with sgControl or sgZNF217. **C,** Representative images (left) and tumor weights (right) of mammary fat pad-implanted tumors formed by MDA-MB-157 cells transduced with sgControl or sgZNF217 (n = 8) and EO771 cells transduced with sgControl or sgZfp217 (n = 10). **D,** Schematic of the experimental workflow for breast cancer patient-derived organoids (PDOs) and associated functional assays. **E–F,** Representative images of RFP and LysoTracker fluorescence (E) and quantification (F) of LysoTracker fluorescence in indicated PDOs transduced with a sgControl or sgZNF217. **G–H,** Representative images of RFP and LiperFluo fluorescence (G) and quantification (H) of LiperFluo fluorescence in indicated PDOs transduced with a sgControl or sgZNF217 following treatment with RSL3 (2 μM, 4 hours). **I–J,** Representative images in brightfield and RFP fluorescence (I) and quantification of organoid mass (J) of indicated PDOs transduced with a sgControl or sgZNF217 after 7 days of culture. **K-L,** Quantification of organoid mass (K) and organoid mean size (L) in sgZNF217-transduced PDOs treated with vehicle or with Lip-1 (5 μM) after 7 days of culture. Each data point represents one PDO. Panel B is shown as mean ± s.d.; n = 3 biologically independent experiments. Tumor weights in C are shown as box plots; the center line indicates the mean, the box limits indicate the first and third quartiles, and whiskers indicate the range. Organoid quantification in F and H is presented as box plots; each data point represents one organoid; the center line indicates the mean, the box limits indicate the first and third quartiles, and whiskers indicate the range. More than four organoids were analyzed for each organoid line. Organoid growth in J is shown as mean ± s.d.; n = 3 biologically independent experiments. Statistics were performed using two-sided unpaired t tests unless otherwise indicated.

For cell line studies, we first disrupted *ZNF217* expression in the human triple-negative breast cancer (TNBC) cell line MDA-MB-157 (**Supplementary Fig. 9A**), which exhibits elevated *ZNF217* copy number (**Supplementary Fig. 9B**), as well as in the murine syngeneic breast cancer cell line EO771 (**Supplementary Fig. 9C**). Loss of *ZNF217* impaired lysosomal acidification and pHrodo-LDL signal in vitro in these models (**Fig. 5B; Supplementary Fig. 9D and 9E**). Notably, orthotopic implantation of MDA-MB-157 and EO771 ZNF217-deficient cells into the mammary fat pad resulted in tumors with reduced weight by 58% and 62%, respectively, compared with tumors arising from isogenic control cells (**Fig. 5C; Supplementary Fig. 9F**).

These findings motivated us to extend our analyses to breast cancer (BC) patient-derived organoids^60^ (PDOs). We evaluated the role of ZNF217 in four independent PDO models: three derived from estrogen receptor-positive (ER+) primary breast cancers (CLO061025, CLO071525-01, and CLO011426) and one derived from a primary TNBC (CLO010626).

Because primary organoids are less amenable to whole population transduction than established cell lines, we cloned either a control sgRNA or one of the previously used sgRNAs targeting human *ZNF217* into a lentiviral backbone expressing RFP, which then allowed us to identify transduced regions by live-cell fluorescence imaging (**Fig. 5D**). We combined this approach with LysoTracker and LiperFluo, two dyes compatible with live imaging that serve as proxies for lysosomal acidity and lipid peroxidation, respectively. Notably, LysoTracker signal was reduced in sgZNF217-RFP-positive regions compared with sgControl-RFP regions (**Fig. 5E and 5F; Supplementary Fig. 10A**) across all four independent PDOs. Moreover, LiperFluo intensity increased in sgZNF217-transduced cells relative to controls following GPX4 inhibition with RSL3 (**Fig. 5G and 5H; Supplementary Fig. 10B**). These results are consistent with the previous data linking lysosomal acidification to enhanced lipoprotein assimilation and ferroptosis resistance across tumor types. Thus, the role of ZNF217 in these phenotypes extends to both breast cancer cell lines and PDOs.

Finally, we assessed the growth and aggressiveness of sgControl- and sgZNF217-transduced PDOs by quantifying two different traits: organoid mass and size. Organoid size was quantified as the projected area of individual organoids, while organoid mass was calculated as the sum of the segmented projected areas of all organoids within each well and was used as a surrogate measure of the total organoid burden. Knocking out *ZNF217* reduced organoid mass and size within a range of 1.6-4.8-fold and 1.8-6.0-fold in all four organoids, respectively (**Fig. 5I and 5J; Supplementary Fig. 10C and 10D**). To determine whether these growth defects were driven by ferroptosis, we treated ZNF217-deficient PDOs with the ferroptosis inhibitor Lip-1. Although the magnitude of rescue varied across the four PDO models, Lip-1 stimulated organoid mass and size in 3 out of the 4 PDOs tested (**Supplementary Fig. 11A and 11B**). When data for each organoid was normalized to vehicle-treated sgZNF217 controls, Lip-1 significantly restored organoid growth across the four independent patient-derived models (**Fig. 5K and 5L**), supporting ferroptosis as a contributor to the impaired growth of ZNF217-deficient breast cancer PDOs.

Together, these results demonstrate that ZNF217 promotes breast cancer PDO growth, in part, through maintenance of lysosomal acidification and resistance to ferroptosis. In summary, we show that ZNF217-mediated transcriptional regulation of *RAB11FIP4* is a key driver of lysosomal acidity, ferroptosis resistance, and tumor aggressiveness. More broadly, these findings indicate that lysosomal acidification is required for ferroptosis evasion, raising the possibility of targeting this axis as a pro-ferroptotic therapeutic strategy.

## DISCUSSION

Lysosome-dependent degradation of extracellular cargo, such as proteins or nucleotides, provides tumors with an important source of nutrients and metabolic flexibility^61^. Recently, several studies have highlighted the role of lipoprotein assimilation, a process similarly dependent on lysosomes, in supporting tumor growth by enabling acquisition of lipid antioxidants that suppress ferroptosis^15,17^. Efficient lipoprotein assimilation requires two coordinated processes: uptake of extracellular lipoproteins followed by delivery to lysosomes and degradation by acidic lipases. Here, we sought to define the mechanisms by which tumor cells actively regulate this latter step of lipoprotein processing. We identify ZNF217 as a transcriptional regulator of this process. By maintaining lysosomal acidification, ZNF217 enables acquisition of lipoprotein-derived antioxidants, which results on tumors resistant to ferroptotic cell death. Importantly, disruption of ZNF217 expression had little effect on lipoprotein uptake itself but instead impaired delivery of internalized cargo to acidic lysosomes. Thus, our findings establish that tumor lipoprotein assimilation is not simply determined by lipoprotein uptake, as we and others have shown previously, but also depends on active maintenance of lysosomal function.

Our investigation of how ZNF217 promotes lysosomal function identifies a molecular pathway through which cancer cells transcriptionally regulate lysosomal acidification. Although lysosomal pH has traditionally been viewed as a constitutive property maintained by the V-ATPase and associated trafficking machinery, less is known about how lysosomal acidification is actively regulated to support the enhanced metabolic demands of tumor cells. We find that ZNF217 enables expression of RAB11FIP4, which in turn maintains RAB7A in its active GTP-bound state, thereby preserving lysosomal acidity. Previous studies have implicated RAB11FIP4 in lysosomal biology^38^, but its precise function remained unclear. Our work identifies the ZNF217–RAB11FIP4 axis as a transcriptionally controlled pathway that maintains lysosomal acidification and demonstrates that impaired RAB7A activation contributes to defective lysosomal function following ZNF217 or RAB11FIP4 loss. Although a role of RAB7A in LDL metabolism has been previously reported^62^, how RAB11FIP4 promotes RAB7A activation remains an important unanswered question. RAB7A has established roles in late endosomal maturation, lysosomal trafficking, and lysosome positioning^63^, processes that can influence lysosomal function and acidity. Future studies are needed to define whether RAB11FIP4 regulates RAB7A activation through control of lysosomal organization, trafficking, or other components of the RAB7A regulatory network.

An unexpected observation from our study is that relatively modest changes in lysosomal pH are sufficient to profoundly alter ferroptosis sensitivity. Loss of RAB11FIP4 increased lysosomal pH by only ∼0.4 units, yet this partial defect enhanced ferroptosis sensitivity and suppressed tumor growth while remaining compatible with basal proliferation in vitro. Importantly, pharmacological restoration of lysosomal acidity using the small molecule C381 rescued these phenotypes. These results suggest that tumors do not require complete disruption of lysosomal function to alter metabolic fitness, but rather depend on maintaining an optimal lysosomal state to support metabolic adaptation.

*ZNF217* is a frequently amplified oncogene in human cancers and has been linked to breast cancer progression^20^ through effects on transcriptional and epigenetic regulation. Here, we identify lysosomal acidification as a key downstream function of this oncogene across established breast cancer cell lines and patient-derived organoids, in addition to other cancer cell lines with high lipoprotein uptake. More broadly, our work expands the emerging relationship between lysosomal function and ferroptosis vulnerability. Previous studies have established lysosomes as important determinants of cellular ferroptosis, as they regulate redox-active iron handling^64,65^, undergo extensive membrane lipid peroxidation upon GPX4 inhibition^64^, and harbor localized antiferroptotic systems^33^. We identify the ZNF217–RAB11FIP4 pathway and maintenance of lysosomal acidity and lipoprotein metabolism as an additional determinant of ferroptosis resistance, highlighting lysosomal manipulation as a potential therapeutic strategy for sensitizing tumors to ferroptotic cell death.

## METHODS

### Compounds

The following compounds were used: ferrostatin-1, liproxstatin-1, ML162, erastin, and RSL3 (Cayman Chemical); puromycin, blasticidin, protease inhibitor cocktail, BCECF-AM, and DMEM (Fisher Scientific); B-27 Supplement, Advanced DMEM/F12, TrypLE Express Enzyme, GlutaMAX Supplement, HEPES, pHrodo Low Density Lipoprotein Conjugate, recombinant human R-Spondin-3, recombinant human KGF (FGF-7), recombinant human Heregulin β1, recombinant human Noggin, recombinant human FGF-10, animal-free recombinant human EGF, and chloroquine (Thermo Scientific); RPMI-1640 and Hanks’ Balanced Salt Solution (HBSS) (Gibco); purified human DiI-LDL and lipoprotein-depleted fetal bovine serum (LPDS) (Kalen Biomedical); T4 ligase, BamHI, NotI, and BsmBI (New England Biolabs); Cultrex BME, Type 3 (R&D Systems); SRI-011381 (C381) (Selleck Chemical); dimethyl sulfoxide (DMSO), Tween-80, polybrene, polyethylene glycol-300 (PEG-300), fetal bovine serum (FBS), DAPI, SB202190, N-acetyl-L-cysteine, nicotinamide, Corning Matrigel Basement Membrane Matrix, and IGEPAL CA-630 (Sigma-Aldrich); TransIT-LT1 transfection reagent (Mirus Bio); adenosine 5′-diphosphate, guanosine-5′-triphosphate, bafilomycin A1, and Hoechst 33342 (MedChemExpress); human NME1 (ProSpec); A83-01 (Tocris); Y-27632 dihydrochloride (AbMole); Primocin (InvivoGen); LiperFluo (Dojindo); LysoTracker Green DND-26 (Cell Signaling Technology); LysoTracker Red DND-99 (Invitrogen); and BODIPY™ 581/591 C11 (Invitrogen).

Antibodies to GAPDH (GTX627408; 1:3000) and β-actin (GTX109639; 1:1000) were from GeneTex; 4-hydroxy-2-nonenal (MHN-020P; 1:200) was from the Japan Institute for the Control of Aging (JaICA); V-ATPase A1 (SC-374475; 1:1000), LAMP2 (SC-18822; 1:2000), and cathepsin C (SC-74590; 1:1000) were from Santa Cruz Biotechnology; citrate synthase (14309S; 1:1000), ZNF217 (82306S; 1:1000), RAB7 (D95F2, 9367S; 1:1000), RAB11 (5589S; 1:1000), phospho-p70 S6K (9234S; 1:1000), p70 S6K (2708S; 1:1000), LAMTOR4 (12284S; 1:1000), LC3B (2775S; 1:1000), Golgin-97 (13192S; 1:1000), HA-tag (3724S; 1:2000), and β-tubulin (2146S; 1:2000) were from Cell Signaling Technology; RAB11FIP4 (HPA021595; 1:1000) and FLAG (F3165; 1:5000) were from Sigma-Aldrich; ATP6V1A (EPR19270; 1:1000) was from Abcam; and Cas9 (NBP2-36440SS; 1:1000) was from Novus Biologicals. Horseradish peroxidase (HRP)-conjugated anti-rabbit (7074S; 1:5000) and anti-mouse (7076S; 1:5000) secondary antibodies were purchased from Cell Signaling Technology. Clean-Blot IP Detection Reagent (21230, 1:5000) was from Thermo Scientific. All antibodies are commercially available and validated by the manufacturer; where indicated, specificity was further confirmed by overexpressing the corresponding cDNA and probing by immunoblot.

### Cell lines and cell culture

All cell lines were purchased from ATCC, authenticated by short tandem repeat profiling, and tested for mycoplasma every 2 months. Cells were maintained at 37°C in 5% CO2 in RPMI-1640 supplemented with 1 mM glutamine, 10% FBS, and penicillin/streptomycin. For lipoprotein-depletion experiments, RPMI-1640 was supplemented with 10% LPDS in place of FBS. Murine breast cancer cell line EO771 was a gift from Srinivas Malladi lab at UT Southwestern Medical Center.

### Cellular lipid peroxidation assays with LiperFluo

Adherent cells were seeded in an 8-well µ-Slide (Ibidi) and treated with 2 μM RSL3 for 2 h, followed by supplementation of the culture medium with 2 μM LiperFluo for 1 h. Signal was collected in the GFP channel using a Zeiss LSM880 confocal microscope, and signal intensity per image was quantified using ZEN 3.9 software.

### C11-BODIPY assay for lipid peroxidation

Cells were seeded in 12-well plates overnight, treated with or without 1 μM ferrostatin-1 (Fer-1) for 2 h, and then treated with RSL3 at the indicated concentrations for 16 h. Cells were trypsinized, resuspended in HBSS containing 2 μM BODIPY™ 581/591 C11, and incubated for 1 h at 37°C. Cells were washed and analyzed by flow cytometry, with signals detected in the FITC (oxidized) and PE (reduced) channels. The relative lipid peroxidation ratio was calculated as FITC/(FITC + PE).

### Generation of knockout and cDNA overexpression cell lines

For lentiviral knockout constructs, annealed oligonucleotides were ligated into lentiCRISPR-v2 or lentiCRISPR-v1-RFP vectors, as indicated, using T4 DNA ligase. For lentiviral knockdown constructs, shRNA sequences targeting CCZ1 were selected from the Genetic Perturbation Platform and cloned into the pLKO-puro vector using T4 DNA ligase.

sgZNF217:

5′-CACCGTGGTGAACGGATCGAGCTG-3′

5′-AAACCAGCTCGATCCGTTCACCAC-3′

sgZfp217:

5′-CACCGGGACGTGGGTTCCTCTCGG-3′

5′-AAACCCGAGAGGAACCCACGTCCC-3′

sgRAB11FIP4:

5′-CACCGCTTGCCGTAGGCCTCGCGG-3′

5′-AAACCCGCGAGGCCTACGGCAAGC-3′

sgRAB11FIP4-H3K4me2-region:

5′-CACCGCGGGCGACAAGTGTCTCCA-3′

5′-AAACTGGAGACACTTGTCGCCCGC-3′

sgRAB11FIP4-H3K27ac-region:

5′-CACCCTATAATGCCAGCGATGCAA-3′

5′-AAACTTGCATCGCTGGCATTATAG-3′

shCCZ1:

5′-CCGGGCTGAGTTTCTTCATCTACAACTCGAGTTGTAGATGAAGAAACTCAGCTTTTTG-3′

5′-AATTCAAAAAGCTGAGTTTCTTCATCTACAACTCGAGTTGTAGATGAAGAAACTCAGC-3′

Guide RNA-resistant ZNF217 and RAB11FIP4 cDNAs, RAB7 wild-type and mutant cDNAs, and RILP-S-tag were synthesized as gene fragments (Twist Bioscience). PCR overlap extension and Gibson assembly were used to clone the cDNAs into the pMXs-IRES-Blast vector. For CRISPR interference (CRISPRi) experiments, the pHR-SFFV-KRAB-dCas9-P2A-mCherry construct was stably introduced into Karpas299 or HEK293T cells.

For virus production, sgRNA, shRNA, or cDNA vectors were cotransfected into HEK293T cells together with packaging vectors (VSV-G and ΔVPR for lentivirus; VSV-G and Gag-Pol for retrovirus) using TransIT-LT1 reagent. Medium was replaced 24 h post-transfection, and virus-containing medium was collected 48 h later and passed through a 0.45-μm filter to remove HEK293T cells. Target cells were seeded 24 h before infection. For viral transduction, infection medium containing 8 μg/mL polybrene was added, and cells were spin-infected at 2,200 rpm for 1.5 h. Fresh medium was added 24 h post-infection, and puromycin (for lentiviral sgRNA and shRNA constructs) or blasticidin (for retroviral overexpression constructs) was added for selection. To establish knockout cell lines, single-cell clones were isolated after puromycin selection and expanded in medium containing 1 μM ferrostatin-1 to increase survival of KOs based on the higher ferroptosis phenotype. Gene knockout was confirmed by immunoblotting.

### Transient transfection of RAB7 cDNAs

HeLa cells were seeded at 200,000 cells per well in 6-well plates. The following day, cells were transfected with 1 μg plasmid (RAB7^WT^, RAB7^T22N^, or RAB7^Q67L^) using TransIT-LT1 reagent. After 24 h, medium was changed to either FBS-containing medium (for LysoTracker staining) or LPDS-containing medium (for pHrodo-LDL staining). After an additional 24 h, cells were stained with the indicated dyes and analyzed by flow cytometry.

### Immunoblotting

Cell pellets were washed with PBS and lysed in buffer containing 10 mM Tris-HCl (pH 7.5), 150 mM NaCl, 1 mM EDTA, 1% Triton X-100, 2% SDS, and 0.1% CHAPS, supplemented with protease inhibitors. Samples were sonicated, incubated on ice for 10 min, and centrifuged at 13,000 × g for 10 min at 4°C. The resulting supernatants were collected, and protein concentrations were measured using the Pierce BCA Protein Assay Kit (Thermo Scientific). Lysates were normalized to 20 μg protein in 20 μL and separated by SDS-PAGE before transfer to Immobilon-P membranes (Millipore). Membranes were blocked with 5% milk in TBS-T and processed using standard immunoblotting procedures.

### Proliferation assays

Cell lines were seeded in triplicate in 96-well plates at 1,000 cells/mL for suspension cells or 500 cells/mL for adherent cells in 0.2 mL medium containing the indicated treatments. Untreated cells were plated in parallel and measured at the initial time point for normalization. After 5 days, 40 μL of CellTiter-Glo reagent (Promega) was added to each well, and luminescence was quantified using an Infinite M Plex plate reader (Tecan). Results are presented as relative fold change (log_2_) in luminescence normalized to the initial cell number.

### Co-immunoprecipitation assays

The human RAB7-binding domain of RILP (RILP-RBD; amino acids 241-320^66^) fused to an S tag was cloned into the pMXs vector. Cells stably expressing HA-tagged RAB7^WT^, RAB7^T22N^, or RAB7^Q67L^ were seeded^67^ in 10-cm culture dishes overnight and transfected with 1 μg of RILP-RBD-S-tag expression plasmid using TransIT-LT1 reagent. The medium was replaced 24 h post-transfection, and cells were collected 72 h post-transfection. Cells were lysed in IP buffer containing 25 mM HEPES, 150 mM NaCl, 5 mM MgCl_2_, 0.5% NP-40, 10% glycerol) supplemented with a protease inhibitor cocktail. Lysates were cleared by centrifugation at 12,000 × g for 15 min at 4°C. Supernatants were incubated with anti-S-tag beads overnight at 4°C with rotation. Beads were collected, washed, and boiled in SDS sample buffer at 95°C for 5 min. Pulldown samples were analyzed by immunoblotting using anti-HA antibody to detect RAB7-HA and anti-S-tag antibody to detect the S-tagged RILP-RBD.

For analysis of V-ATPase V0-V1 assembly^68^, cells were lysed in IP buffer containing 50 mM HEPES (pH 7.4), 150 mM KCl, 1 mM MgCl_2_, 1 mM EGTA, 1 mM DTT, 10% glycerol, and 0.3% digitonin supplemented with protease inhibitor cocktail. Lysates were cleared by centrifugation, and the resulting supernatants were incubated with anti-ATP6V0A1 antibody overnight at 4°C with rotation. Protein A/G magnetic beads were then added, and samples were incubated for an additional 2 h at 4°C. Beads were collected, washed, and boiled in SDS sample buffer at 95°C for 5 min. Immunoprecipitated proteins were analyzed by immunoblotting using antibodies against ATP6V1A and ATP6V0A1.

### CRISPR-based genetic screens

The human epigenetic and transcriptional regulator sgRNA library was generated in previous work^23^. The histone mark CUT&RUN and Lyso-IP sgRNA libraries were curated from the corresponding datasets generated in this study. The CUT&RUN library comprised 366 candidate genes selected primarily from differential H3K4me2 and H3K4me3 analyses (|log₂FC| ≥ 2 and FDR<0.05 for H3K4me2; |log₂FC| ≥ 1.3 and FDR < 0.05 for H3K4me3; *Supplementary Table 1*), together with transcription factors identified by H3K4me2 motif enrichment. Oligonucleotides (Twist Bioscience) were annealed and cloned into lentiCRISPR-v2 using a Golden Gate reaction. These lentiviral libraries were transfected into HEK293T cells to generate viral supernatant as described above. Cells were infected at a multiplicity of infection of 0.7 and selected with puromycin. For the screens, 30 million cells (human epigenetic library) or 3 million cells (focused, smaller libraries) were infected and cultured for 14 population doublings under sublethal RSL3 (70 nM). After culture, 30 million cells were harvested and genomic DNA was extracted using a DNeasy Blood and Tissue Kit (Qiagen).

For flow cytometry-based pHrodo-LDL screens, cells were transduced, selected, and expanded as described above. An initial pool of 30 or 3 million cells was collected and placed in RPMI with LPDS overnight before sorting. Two hours prior to sorting, cells were incubated with 10 μg/mL pHrodo-LDL in HBSS. Cells were then washed, resuspended in HBSS with DAPI, and passed through a 40-μm strainer. Sorting was performed on a FACSAria III (BD Biosciences), collecting the top and bottom 10% of pHrodo-LDL-fluorescing cells. Sorted cells were maintained in RPMI with ferrostatin-1 (1 μM) until 30 or 3 million cells were collected for sgRNA analysis.

sgRNA inserts were PCR-amplified using condition-specific primers, purified, and sequenced on a NextSeq 2000 (Illumina). Sequencing reads were mapped, and the abundance of each sgRNA was quantified. Gene scores were defined as the median log2 fold change in abundance of all sgRNAs targeting a given gene between conditions; a gene score below −1 was considered significant. Screens were analyzed using Python (v2.7.13), R (v3.3.2), and Unix (v4.10.0-37-generic-x86_64).

Gene scores from the epigenetic and transcriptional regulator, CUT&RUN-focused, and Lysosome-focused CRISPR screens are included in *Supplementary Tables 4–6*, respectively.

### Subcutaneous or orthotopic xenograft tumors in mice

All animal studies were approved by the Institutional Animal Care and Use Committee (IACUC) at the University of Texas Southwestern Medical Center and were conducted in accordance with institutional guidelines. C57BL/6J (JAX #000664) and NSG (JAX #005557) mice were obtained from The Jackson Laboratory. Mice were housed in a pathogen-free environment at 68–79°F and 30–70% humidity, with a 12-h light/dark cycle, and fed standard chow ad libitum unless otherwise noted. The IACUC-approved tumor size limit of 2 cm in diameter or 10% of body weight was not exceeded in any experiment. Sample sizes were not predetermined by statistical power calculations but were based on prior experience with these assays. For pharmacological treatment experiments, mice were randomly allocated to treatment groups, and investigators were blinded to treatment during analysis.

Xenograft tumors were initiated by injecting the following numbers of cells in DMEM with 30% Cultrex: 10,000 cells in 100 μL for Karpas299 parental and knockout lines, subcutaneously; 50,000 cells in 100 μL for HeLa parental and knockout lines, subcutaneously;

50,000 cells in 100 μL for NCI-H838 parental and knockout lines, subcutaneously;

100,000 cells in 100 μL for PaTu 8988t parental and knockout lines, subcutaneously;

200,000 cells in 50 μL for PaTu 8988t parental and knockout lines, orthotopically;

500,000 cells in 50 μL for MDA-MB-157 parental and knockout lines, orthotopically;

100,000 cells in 50 μL for EO771 parental and knockout lines, orthotopically.

Experiments using EO771 cells were performed in immunocompetent C57BL/6J mice, whereas all other tumor studies were conducted in immunodeficient NSG mice. For subcutaneous tumor studies, cells were implanted into the bilateral flanks of 6–14-week-old male or female mice. For experiments comparing knockout tumor and isogenic control growth, knockout cells and isogenic controls were implanted into opposite flanks of the same mouse. For pharmacological studies, mice were implanted bilaterally with tumors of a single genotype, with one tumor established in each flank, and were randomized to receive vehicle, C381, or Lip-1 as indicated. For orthotopic tumor studies, PaTu 8988t cells were injected into the pancreas using a 29-gauge insulin syringe (BD), whereas MDA-MB-157 and EO771 cells were injected into the fourth mammary fat pad of female mice.

The anti-ferroptotic compound liproxstatin-1 (Lip-1) was reconstituted in DMSO and mixed with PEG-300, Tween-80, and water to a final solvent composition of 5.1% DMSO, 40% PEG-300, 2% Tween-80, and 52.9% water. The solution was vortexed, sonicated, centrifuged, and stored in aliquots at −70°C; a vehicle solution without Lip-1 was prepared identically for comparison. Lip-1 (10 mg/kg) or vehicle was administered by intraperitoneal injection starting 3 days before tumor implantation and every other day thereafter until the end of the experiment.

The lysosomal agonist SRI-011381 (C381) was reconstituted in DMSO and mixed with PEG-300, Tween-80, and water to a final solvent composition of 5% DMSO, 40% PEG-300, 5% Tween-80, and 50% water, as previously described^56^. The solution was vortexed, sonicated, centrifuged, and stored in aliquots at −70°C; a vehicle solution without C381 was prepared identically for comparison. C381 (30 mg/kg) or vehicle was administered by intraperitoneal injection twice weekly until the end of the experiment.

### Immunofluorescence assays

Cells were seeded onto glass coverslips the day before image acquisition, fixed with 4% paraformaldehyde for 10 min, permeabilized with 0.3% Triton X-100 for 10 min, and blocked with 5% BSA for 1 h. Cells were incubated with primary antibody overnight at 4°C, followed by fluorophore-conjugated secondary antibody for 1 h at room temperature. DAPI was used as a counterstain, and slides were mounted with VECTASHIELD antifade mounting medium (H-1900, Vector Laboratories).

### Live-cell imaging of pHrodo-LDL and DiI-LDL assimilation

Cells were seeded onto 8-well µ-Slides (Ibidi) in LPDS-containing medium overnight, then incubated with 10 μg/mL pHrodo-LDL or DiI-LDL prior to live imaging. Hoechst 33342 was used as a nuclear counterstain, and CellMask was used to stain the cell membrane. Cells were imaged with a Zeiss LSM880 confocal microscope, and signal intensity was quantified using ZEN 3.9 software.

### Flow cytometry determination of cellular assimilation of pHrodo-LDL and DiI-LDL

Adherent and suspension cell lines were plated in six-well plates at 150,000 or 300,000 cells per well, respectively, in RPMI with LPDS. After 24 h, medium was removed and replaced with fresh medium containing 10 μg/mL DiI-LDL or pHrodo-LDL, and cells were incubated for 1 h. Cells were collected, washed twice with cold HBSS, resuspended in HBSS with DAPI, passed through a 40-μm strainer, and kept on ice until analysis. Data were acquired on a FACSCanto RUO (BD Biosciences), collecting PE signal for DiI-LDL, or GFP signal for pHrodo-LDL, to assess uptake/assimilation via median fluorescence intensity (MFI) shift. At least 5,000 events were recorded per sample, and data were analyzed using FlowJo v10.

### LysoTracker staining

For flow cytometric analysis, cells were incubated with LysoTracker Green DND-26 (1:10,000) for 10 min at 37°C, harvested, and resuspended in HBSS containing DAPI prior to acquisition on a FACSCanto RUO (GFP channel). For live-cell imaging, cells were seeded in 8-well µ-Slides and stained with LysoTracker Green DND-26 (1:10,000), CellMask Deep Red (1:1,000), and Hoechst 33342 for 10 min at 37°C, followed by imaging on a Zeiss LSM880 confocal microscope. Unless otherwise indicated, images were acquired at a single optimal focal plane selected based on the CellMask signal to ensure accurate cell boundary delineation. Cell boundaries were segmented using the CellMask channel, and LysoTracker fluorescence was quantified within each segmented cell. For validation of the image analysis strategy (Supplementary Fig. 6B), live-cell images of HeLa *RAB11FIP4*_KO cells and isogenic controls were additionally acquired as z-stacks using an Agilent BioTek Cytation C10 confocal imaging reader. The focal plane with the strongest CellMask signal was selected for analysis using the same CellMask-based segmentation and LysoTracker fluorescence quantification pipeline.

### Genetically encoded lysosomal pH reporters

Lysosomal pH was independently assessed using the ratiometric genetic reporters pHlare^44^ (164477, Addgene) and RpH^45^ (225135, Addgene). Cells were transiently transfected with each construct and, 72 h post-transfection, seeded into an 8-well µ-Slide for confocal imaging. Relative lysosomal pH was calculated as the ratio of FITC to mCherry fluorescence intensity.

### Lysosomal pH standard curve

A series of calibration buffers spanning pH 5.0–7.0 (0.5-unit increments) was prepared. Cells were treated with the H⁺/K⁺ ionophore nigericin to equilibrate intracellular and extracellular pH, stained with LysoSensor Yellow/Blue DND-160^69^, and resuspended in the corresponding calibration buffers. Yellow (BV510) and blue (DyeCycle Violet) fluorescence signals were acquired by flow cytometry, and lysosomal pH was determined from the yellow-to-blue fluorescence ratio referenced against the standard curve.

### Cytosolic pH measurement

Cytosolic pH was measured using the ratiometric pH-sensitive dye BCECF-AM. Cells were loaded with BCECF-AM and analyzed by flow cytometry, as previously described^70^.

### Ex vivo tumor LysoTracker staining

Tumors were harvested and immediately placed in ice-cold dissection buffer (15 mM HEPES, 6.5 mg/mL glucose, 1.3 mM MgSO4, 20 mM KCl, 1% penicillin/streptomycin in 1× HBSS). Tumors were embedded in 3% low-melting-point agarose (Invitrogen, 16520-100) on ice and sectioned into 200 μm slices using a vibratome (Leica). Sections were transferred into dissection buffer-containing 8-well µ-Slides, stained with LysoTracker Green DND-26 for 30 min, and counterstained with Hoechst 33342. Live sections were then imaged by confocal microscopy.

### Measurement of RAB7-GTP levels

RAB7-HA was immunoprecipitated using anti-HA magnetic beads. Following washes, nucleotides were eluted by resuspending the IP beads in 50 μL nucleotide elution buffer (25 mM HEPES pH 7.4, 100 mM NaCl, 5 mM EDTA) and heating at 60°C for 5 min. Beads were briefly spun down, and the supernatant containing eluted nucleotides was collected and kept on ice or stored at −80°C until use. In parallel, the remaining IP beads were resuspended in 2× SDS loading buffer and heated at 95°C for 5 min, and the resulting supernatant was collected for western blot analysis of HA, alongside input samples, to normalize for RAB7-HA pulldown efficiency.

We followed a previously described protocol^71^. Briefly, GTP bound to immunoprecipitated RAB7 was quantified using an NME1-coupled luminescence assay. Recombinant NME1 (nucleoside diphosphate kinase; specific activity 1200 U/mg) was diluted 1,000-fold in reaction buffer (50 mM HEPES pH 7.4, 150 mM NaCl, 10 mM MgCl2, 1 mM DTT, 0.05% Tween-20, 0.1% BSA) to a working concentration of 1.2 mU/μL. A 2× Conversion Mix containing 1 mM ADP and 0.01 U/mL NME1 was prepared in reaction buffer. In a 96-well plate compatible with CellTiter-Glo (CTG) luminescence detection, 20 μL of nucleotide eluate was combined with 20 μL of 2× Conversion Mix and incubated overnight at room temperature to allow NME1-catalyzed transfer of the γ-phosphate from GTP to ADP, generating ATP in stoichiometric proportion to the amount of GTP present in the sample. Following incubation, 40 μL of CellTiter-Glo reagent was added to each well, and plates were incubated for 10 min in the dark before luminescence was measured.

A standard curve was generated in parallel by preparing serial dilutions of purified GTP (0, 125, 250, 500, and 1000 nM) in nucleotide elution buffer and subjecting them to the same conversion and CellTiter-Glo detection steps. Luminescence values were background-subtracted using the 0 nM GTP control, and GTP concentrations in experimental samples were interpolated from this standard curve and normalized to RAB7-HA levels determined by densitometric analysis of the corresponding western blots.

### Histology and immunohistochemistry

Xenograft tissues were fixed in 10% formalin, embedded in paraffin, and sectioned into 4 μm slices. Following deparaffinization and rehydration, antigen retrieval was performed for ZNF217 staining using AR Citra Plus Solution in a pressure cooker but was omitted for 4-HNE staining. Sections then underwent endogenous peroxidase blocking, followed by blocking with normal goat serum for 20 min, then incubated with anti-ZNF217 or 4-HNE antibody (0.5 μg/mL) for 1 h, followed by biotinylated secondary antibody for 30 min and VECTASTAIN ABC reagent (PK-4000, Vector Laboratories). Sections were developed in DAB peroxidase substrate (SK-4100, Vector Laboratories) for 10 min and counterstained with hematoxylin (H-3401, Vector Laboratories). Sections were mounted using DPX Mountant for histology (06522, Sigma-Aldrich). Slides were scanned using a NanoZoomer S60 (Hamamatsu Photonics) and analyzed using NDP.view2 v2.9.29 at 20× magnification. Intensity of 4-HNE signal across the entire tissue section was quantified using ImageJ (v1.54g): images were subjected to color deconvolution (H DAB method) to isolate the DAB channel, converted to 8-bit grayscale, and inverted; regions of interest encompassing the whole tissue section were selected, and mean DAB intensity was measured as a readout of 4-HNE staining.

### CUT&RUN analysis of epigenetic alterations in *ZNF217* knockout cells

CUT&RUN was performed using the CUTANA ChIC/CUT&RUN Kit (v4; EpiCypher) and CUT&RUN Library Prep Kit following the manufacturer’s protocol. Briefly, 500,000 Karpas299 control or ZNF217 knockout (ZNF217-KO) cells were immobilized on Concanavalin A-coated beads and incubated overnight with antibodies against H3K4me2, H3K4me3 or H3K27ac, with IgG as a negative control. Chromatin was digested with pAG-MNase, and released DNA fragments were collected in Stop Master Mix supplemented with E. coli spike-in DNA. Libraries were amplified for 14 cycles and sequenced on an Illumina NextSeq platform (paired-end, 150 bp). Reads were aligned using Bowtie2, and peaks were called using MACS2 (n = 3 biological replicates).

### Immunoprecipitation and LC-MS determination of ZNF217 binding partners

Karpas299 ZNF217 knockout cells transduced with either empty vector (EV) or 3×FLAG-ZNF217 were lysed in IP lysis buffer supplemented with protease inhibitors. Equal amounts of protein lysate were incubated with anti-FLAG M2 magnetic beads (Sigma-Aldrich) overnight at 4°C. Following washes, bound proteins were eluted in SDS sample buffer by boiling at 95°C for 5 min, separated on an 8% SDS-PAGE gel, and visualized by Coomassie staining. Gel regions containing the stacked protein bands were excised, digested with trypsin, and analyzed by LC-MS/MS using an Orbitrap Fusion Lumos mass spectrometer coupled to an UltiMate 3000 nano-HPLC with a 75 μm × 75 cm column and a 90-min gradient. Raw data were searched against the reviewed human UniProt database using Proteome Discoverer 3.0. ZNF217-associated proteins were identified based on enrichment in 3×FLAG-ZNF217 samples relative to EV controls.

### Immunopurification of lysosomes (Lyso-IP)

We followed a previously described protocol^40^. Karpas299 cells stably expressing TMEM192-3×HA (HA-Lyso cells) or TMEM192-2×FLAG (Control-Lyso cells) were collected and lysed in KPBS using a Dounce tissue grinder (Kimble). Cell debris was removed by centrifugation at 1,000 × g for 90 s. Anti-HA magnetic beads (88837, Thermo Scientific) were used to pull down lysosomes. Purified lysosomes were lysed in 80 μL lysis buffer, with 10 μL used for protein quantification and the remainder used for western blotting following boiling in Laemmli buffer. The protein gel was stained with Coomassie Brilliant Blue solution (1610436, Bio-Rad) and excised for proteomic analysis.

### RNA extraction and RNA sequencing

RNA was extracted using TRIzol (Thermo Fisher Scientific) and purified using the RNeasy Mini Kit (Qiagen). RNA concentration was measured using a Qubit fluorometer and the Qubit RNA High Sensitivity Kit (Invitrogen).

RNA-seq libraries were prepared using the NEBNext Ultra II Directional RNA Kit together with the NEBNext Poly(A) mRNA Isolation Module (New England Biolabs) according to the manufacturer’s instructions and indexed using standard NEB indices. Sequencing reads were trimmed to remove adapter sequences and low-quality bases (quality score < 25), and reads shorter than 35 bp were discarded.

Reads were aligned to the GRCh38 reference genome using HISAT2 v2.2.1, and duplicate reads were marked using SAMBAMBA v1.0.1. Gene, transcript, and exon counts were generated using featureCounts v2.1.1. Differential gene expression analysis was performed using edgeR v3.21 and DESeq2.

### Real-time quantitative PCR

Cells were collected, washed once with PBS, and RNA was isolated as described above. iScript Reverse Transcription Supermix (1708841, Bio-Rad) was used to generate cDNA. Quantitative PCR was performed using SYBR Green Mastermix (NC2203420, ABclonal Technology), with TBP or HPRT1 used as internal controls for human genes and Actb used as the internal control for mouse genes. Primer sequences were as follows:

RAB11FIP4_For: CCTGCTCAATGACTTGGAAGCC

RAB11FIP4_Rev: ACAGGTCTTCGGTGCTGCCATT

CCZ1_For: TTGCCGAAGACTGGACAGCATC

CCZ1_Rev: TGTGCTCTTCTCGGCGAGATTC

TBP_For: TGTATCCACAGTGAATCTTGGTTG

TBP_Rev: GGTTCGTGGCTCTCTTATCCTC

HPRT1_For: CATTATGCTGAGGATTTGGAAAGG

HPRT1_Rev: CTTGAGCACACAGAGGGCTACA

Mouse Zfp217_For: AGACCTCAGCACCACTCTGGAA

Mouse Zfp217_Rev: GGCTTCGCTTTTCCTCCATCATC

Mouse Actb_For: GTGACGTTGACATCCGTAAAGA

Mouse Actb_Rev: GCCGGACTCATCGTACTCC

### Patient-derived organoid culture

Organoids were dissociated with TrypLE for 5 min into small cell clusters and resuspended in Matrigel Basement Membrane Matrix in 24-well plates. After the Matrigel dome solidified, organoid culture medium was added to each well, and cultures were maintained at 37°C in 5% CO2. The organoid culture medium consisted of 250 ng/mL R-Spondin-3, 5 nM Heregulin β1, 5 ng/mL FGF-7, 20 ng/mL FGF-10, 100 ng/mL Noggin, 5 ng/mL EGF, 0.5 μM A83-01, 5 μM Y-27632, 0.5 μM SB202190, 1.25 mM N-acetylcysteine, 5 mM nicotinamide, 10 mM HEPES, GlutaMAX, Primocin, penicillin/streptomycin, B-27 supplement, and Advanced DMEM/F12, as previously described^60,72^.

A summary of the clinical characteristics of the breast cancer patient-derived organoids are provided in *Supplementary Table 7*.

### Lentiviral transduction of organoids

Organoids were dissociated with TrypLE for 5 min and resuspended in 1 mL organoid medium. Lentivirus (produced in DMEM containing 10% FBS) was added at a 1:1 volume ratio relative to the organoid suspension, together with polybrene (final concentration 10 μg/mL), and the mixture was transferred to a 6-well ultra-low attachment plate. Organoids were spin-infected by centrifugation at 600 × g for 1 h at room temperature, followed by overnight incubation at 37°C, after which media with lentiviral particles was replaced by fresh media.

### Organoid growth assay

To compare growth between sgControl- and sgZNF217-transduced organoids, virus-transduced organoids were embedded in Matrigel and cultured for 7 days. Because organoids grown in Matrigel domes occupy multiple focal planes, precluding accurate imaging, domes were briefly dissociated with TrypLE for 3 min to recover intact organoids, which were then seeded into 24-well ultra-low attachment plates (3473, Corning) to settle into a single focal plane for imaging.

Organoid mass and average organoid size were quantified using ImageJ (v1.54g). Images were first processed by automatic brightness/contrast adjustment and background subtraction (rolling ball radius, 40 pixels). Organoids were then segmented using the Default threshold algorithm, and particle analysis was performed with a size threshold of 50 pixels to infinity and a circularity range of 0.0-1.0. Total organoid area (organoid mass) and average organoid size were calculated from the segmented particles.

### Live imaging of LysoTracker and LiperFluo in organoids

Transduced organoids were embedded in Matrigel domes, seeded onto 8-well µ-Slides (80807, Ibidi), and cultured overnight. For LysoTracker imaging, organoids were incubated with LysoTracker Green DND-26 (1:10,000) for 15 min and washed once with HBSS prior to live imaging in the FITC channel. For LiperFluo imaging, organoids were treated with 2 μM RSL3 for 4 h, followed by incubation with 2 μM LiperFluo for 1 h, then washed once with HBSS and imaged live.

### DepMap analysis

ZNF217 copy number data were obtained from the Cancer Dependency Map^73^ (DepMap, Broad Institute). Gene-level copy number values were extracted from all annotated cancer cell lines, grouped by tissue lineage, transformed as log_2_(copy number + 1), and visualized using GraphPad Prism.

### Statistics and reproducibility

Statistical analyses were performed using GraphPad Prism v10.2.1 and Microsoft Excel v15.21.1. Error bars, P values, and statistical tests are reported in the corresponding figure legends. Data were considered statistically significant at P < 0.05. In vitro experiments were repeated at least three times, and mouse experiments were repeated at least twice, with consistent results across both technical and biological replicates. Schematics and figures were created using Adobe Illustrator v29.5.

## DATA AVAILABILITY

The data and reagents generated in this study are available upon request from the corresponding author Javier Garcia-Bermudez.

## AUTHOR’S CONTRIBUTIONS

**R. Wu:** Conceptualization, data curation, formal analysis, investigation, methodology, visualization, writing – review and editing. **S.-C. Hsu:** Data curation, formal analysis, investigation, methodology, writing – review and editing. **L. Sang:** Investigation, methodology, writing – review and editing. **M. Yu:** Investigation, methodology. **Y.J. Kim:** Methodology. **M. Choe:** Methodology. **C. Hauer:** Resources. **L. Cai:** Formal analysis. **A.B. Hanker:** Resources. **I. S. Chan:** Resources. **H.R. Shin:** Methodology, resources, writing – review. **J. Garcia-Bermudez:** Conceptualization, supervision, funding acquisition, writing – original draft, project administration, writing – review and editing.

## ACKNOWLEDGMENTS

We thank members of the Garcia-Bermudez laboratory for input. J.G.B. is supported by the NIH (1R01CA316462-01, 1R01DK145539-01, DP2GM159178), the Cancer Prevention and Research Institute of Texas (CPRIT RR210059, RP260791), and the American Cancer Society (ACS RSG-24-1255384-01), and is a Pew-Stewart Cancer Scholar. S.-C.H. is funded by a Human Frontiers Postdoctoral Fellowship (HFSP-LT0006/2024-L). L.S. is supported by an American Cancer Society Postdoctoral Fellowship (PF-25-1301347-01-PFMBB), and M.Y. by an American Heart Association predoctoral fellowship (26PRE1551678). I.C. is supported by Susan G. Komen (CCR231010879), and the NIH (1K08CA270188-01A1). L.C. is supported by NIH (R01CA285336) and CPRIT (RP250561). HRS is supported by the NIH (R35GM165590), the Cancer Prevention and Research Institute of Texas (CPRIT RP250535), the Mary Crowley Cancer Research Foundation, and is a Searle Scholar and Pew Biomedical Scholar. We thank the UTSW Medical Center Whole Brain Microscopy Facility and Dr. Denise Ramirez (NIH S10 awards 1S10OD032267-01). We acknowledge the assistance of the Cancer Organoid Innovation Lab (COIL) at UT Southwestern Simmons Cancer Center, which is supported in part by the National Cancer Institute under award number P30 CA142543.

## SUPPLEMENTARY FIGURE LEGENDS

**Supplementary Figure 1.**
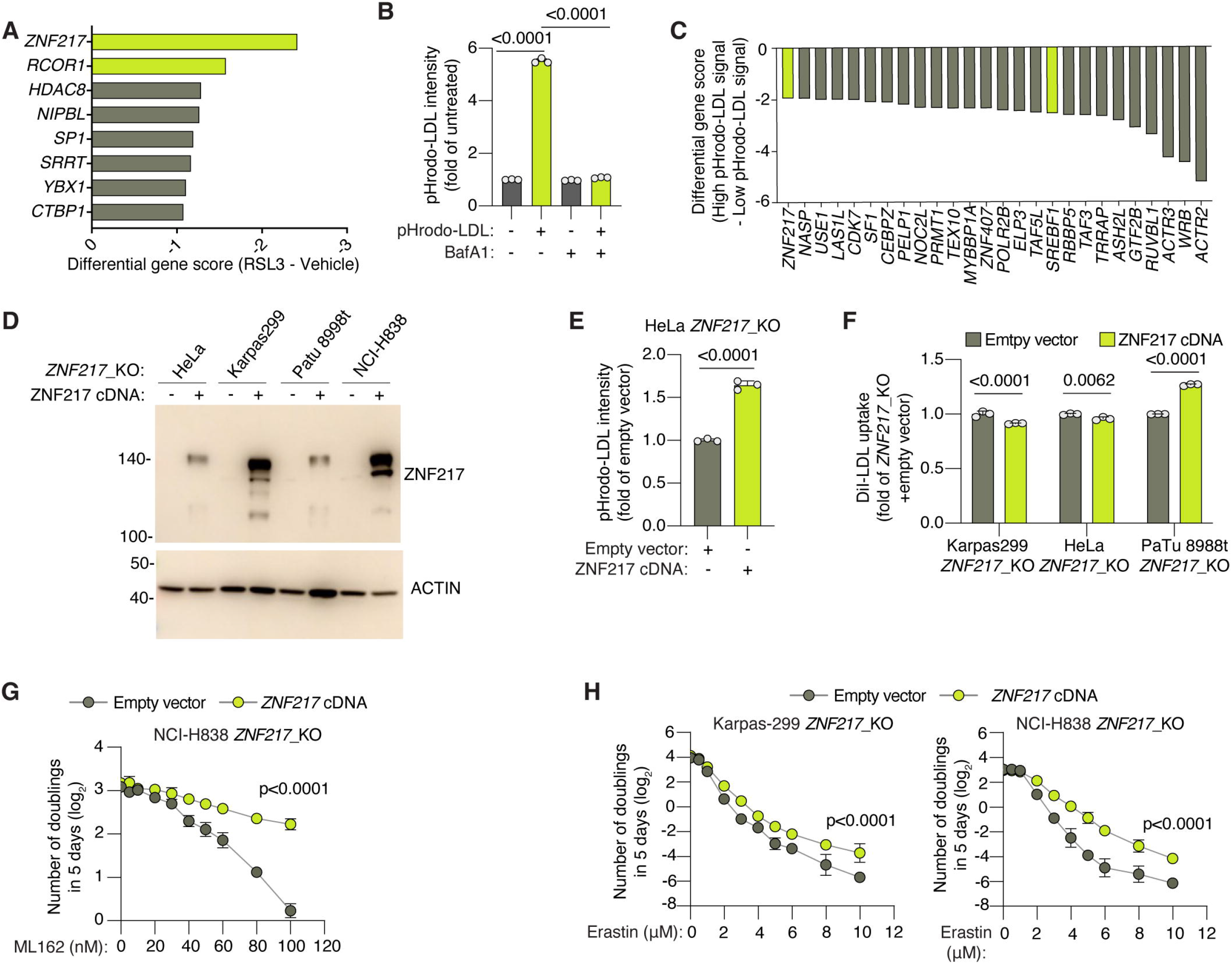
ZNF217 regulates lysosomal lipoprotein assimilation and ferroptosis resistance in cancer cells. **A,** Differential gene scores for the highest-ranked hits identified in the RSL3-treatment proliferation CRISPR screen in Karpas299 cells. **B,** Flow cytometry analysis of pHrodo-LDL fluorescence in cells treated with or without bafilomycin A1 (BafA1, 100 nM). **C,** Differential gene scores of the highest-ranked hits identified in the flow cytometry-based pHrodo-LDL CRISPR screen in PaTu 8988t cells. **D,** Immunoblot analysis of ZNF217 in the indicated *ZNF217*_KO cell lines expressing ZNF217 cDNA or an empty vector. ACTIN is the loading control. **E,** Flow cytometry analysis of pHrodo-LDL fluorescence in HeLa *ZNF217*_KO cells expressing an empty vector or ZNF217 cDNA. **F,** Flow cytometry analysis of DiI-LDL uptake in the indicated *ZNF217*_KO cell lines expressing an empty vector or ZNF217 cDNA. **G,** Number of doublings (log_2_) in 5 days of NCI-H838 *ZNF217*_KO cells expressing an empty vector or ZNF217 cDNA after treatment with the indicated concentrations of ML162. **H,** Number of doublings (log_2_) in 5 days of Karpas299 *ZNF217*_KO (left) and NCI-H838 *ZNF217*_KO cells (right) expressing an empty vector or ZNF217 cDNA after treatment with the indicated concentrations of erastin. Data in B and E–H are mean ± s.d.; n = 3 biologically independent experiments. Panel D is a representative immunoblot. Statistics were performed using two-sided unpaired t tests unless otherwise indicated. Data in B, G, and H were analyzed by two-way ANOVA followed by Šídák’s multiple-comparison test.

**Supplementary Figure 2.**
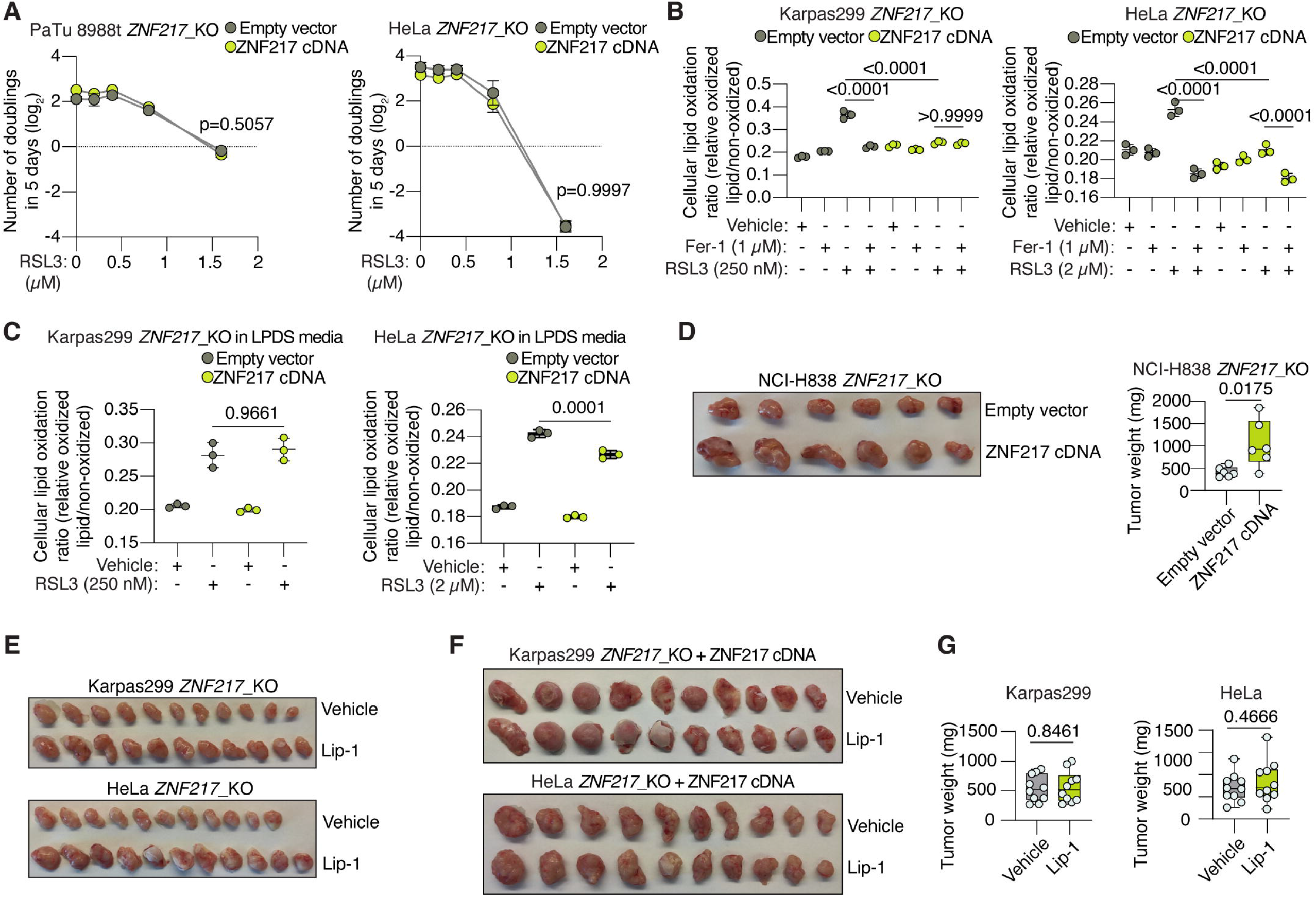
ZNF217 protects cancer cells from ferroptosis in a lipoprotein-dependent manner. **A,** Number of doublings (log_2_) in 5 days of PaTu 8988t (left) and HeLa (right) *ZNF217*_KO cells expressing an empty vector or ZNF217 cDNA after treatment with the indicated RSL3 concentrations in lipoprotein-depleted medium (LPDS). **B,** Quantification of lipid oxidation by BODIPY-C11 fluorescence in Karpas299 (left) and HeLa (right) *ZNF217*_KO cells expressing an empty vector or ZNF217 cDNA after treatment with the indicated concentrations of RSL3 in the presence or absence of ferrostatin-1 (Fer-1; 1 μM). **C,** Quantification of lipid oxidation by BODIPY-C11 fluorescence in Karpas299 (left) and HeLa (right) *ZNF217*_KO cells expressing an empty vector or ZNF217 cDNA cultured in lipoprotein-depleted medium (LPDS) and after treatment with vehicle or the indicated concentrations of RSL3 for 16 h. **D,** Representative images (left) and tumor weights (right) of tumors formed by NCI-H838 *ZNF217*_KO cells expressing an empty vector or ZNF217 cDNA (n = 6). **E,** Representative images of tumors formed by Karpas299 or HeLa *ZNF217*_KO cells after in vivo supplementation with vehicle or Lip-1. **F–G,** Representative images (F) and tumor weights (G) formed by ZNF217-expressing Karpas299 or HeLa cells after in vivo supplementation with vehicle or Lip-1 (n = 10 tumors per treatment group). In vitro data (A–C) are mean ± s.d.; n = 3 biologically independent experiments. For box plots, the center line indicates the median, the box limits indicate the first and third quartiles, and whiskers indicate the range. Statistics were performed using two-sided unpaired t tests unless otherwise indicated. Data in A–C were analyzed by two-way ANOVA followed by Šídák’s multiple-comparison test.

**Supplementary Figure 3.**
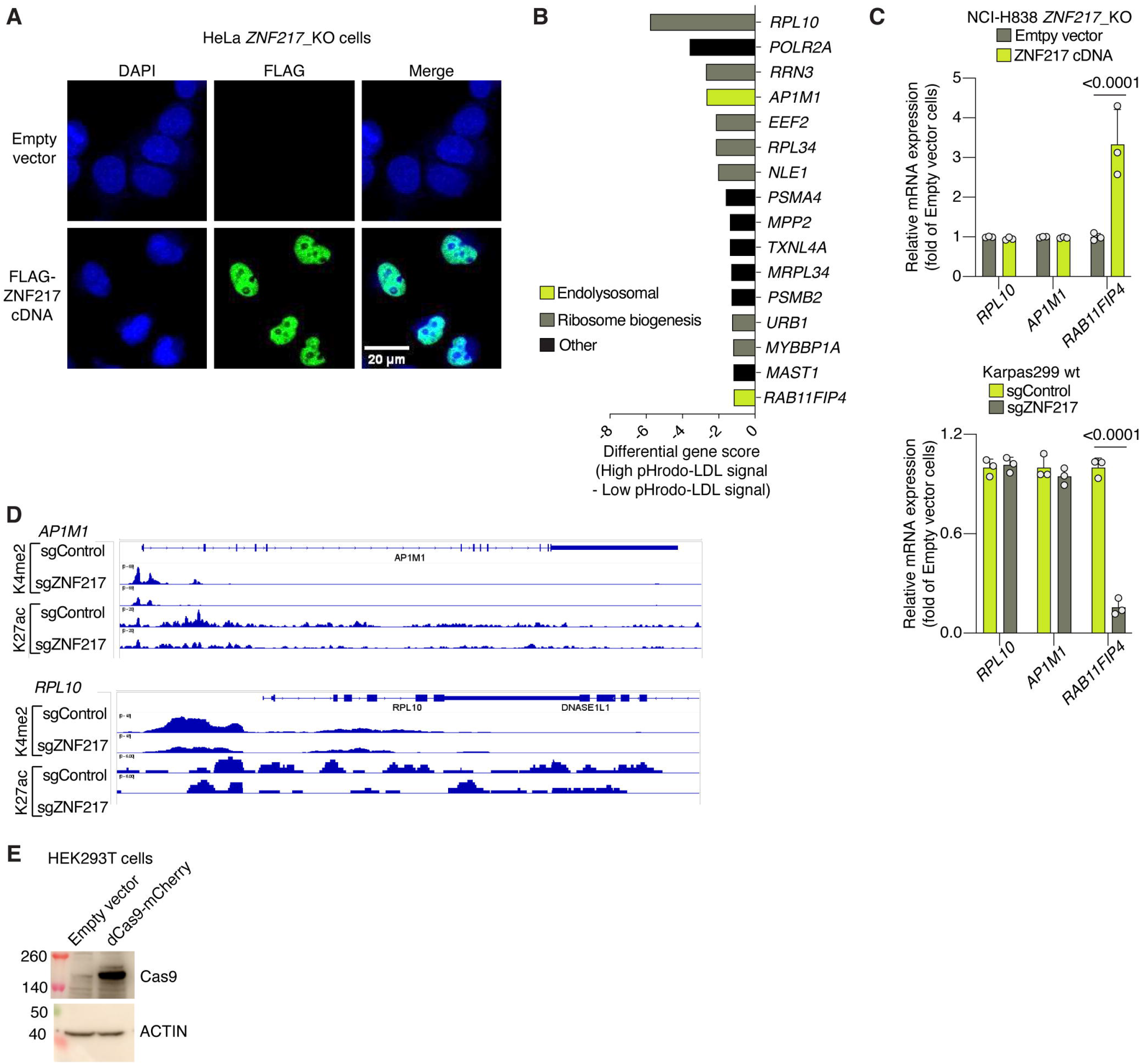
ZNF217 promotes RAB11FIP4 expression. **A,** Representative immunofluorescence images of ZNF217 and DAPI in HeLa *ZNF217*_KO cells expressing an empty vector or FLAG-ZNF217 cDNA. Scale bar, 20 μm. **B,** Differential gene scores from the pHrodo-LDL CRISPR screen in NCI-H838 cells using a focused library of genes identified in CUT&RUN experiments. **C,** Relative mRNA expression of *RPL10*, *AP1M1*, and *RAB11FIP4* in NCI-H838 *ZNF217*_KO cells expressing an empty vector or ZNF217 cDNA (left), and in Karpas299 cells transduced with sgControl or sgZNF217 (right). **D,** H3K4me2 and H3K27ac CUT&RUN profiles at the *AP1M1* and *RPL10* loci in Karpas299 cells expressing sgControl or sgZNF217. **E,** Immunoblot analysis of dCas9 in HEK293T cells expressing dCas9-mCherry or empty vector. ACTIN serves as a loading control. Data in C are mean ± s.d.; n = 3 biologically independent experiments. Panel D is a representative CUT&RUN track; panel E is a representative immunoblot. Statistics were performed using two-sided unpaired t tests unless otherwise indicated.

**Supplementary Figure 4.**
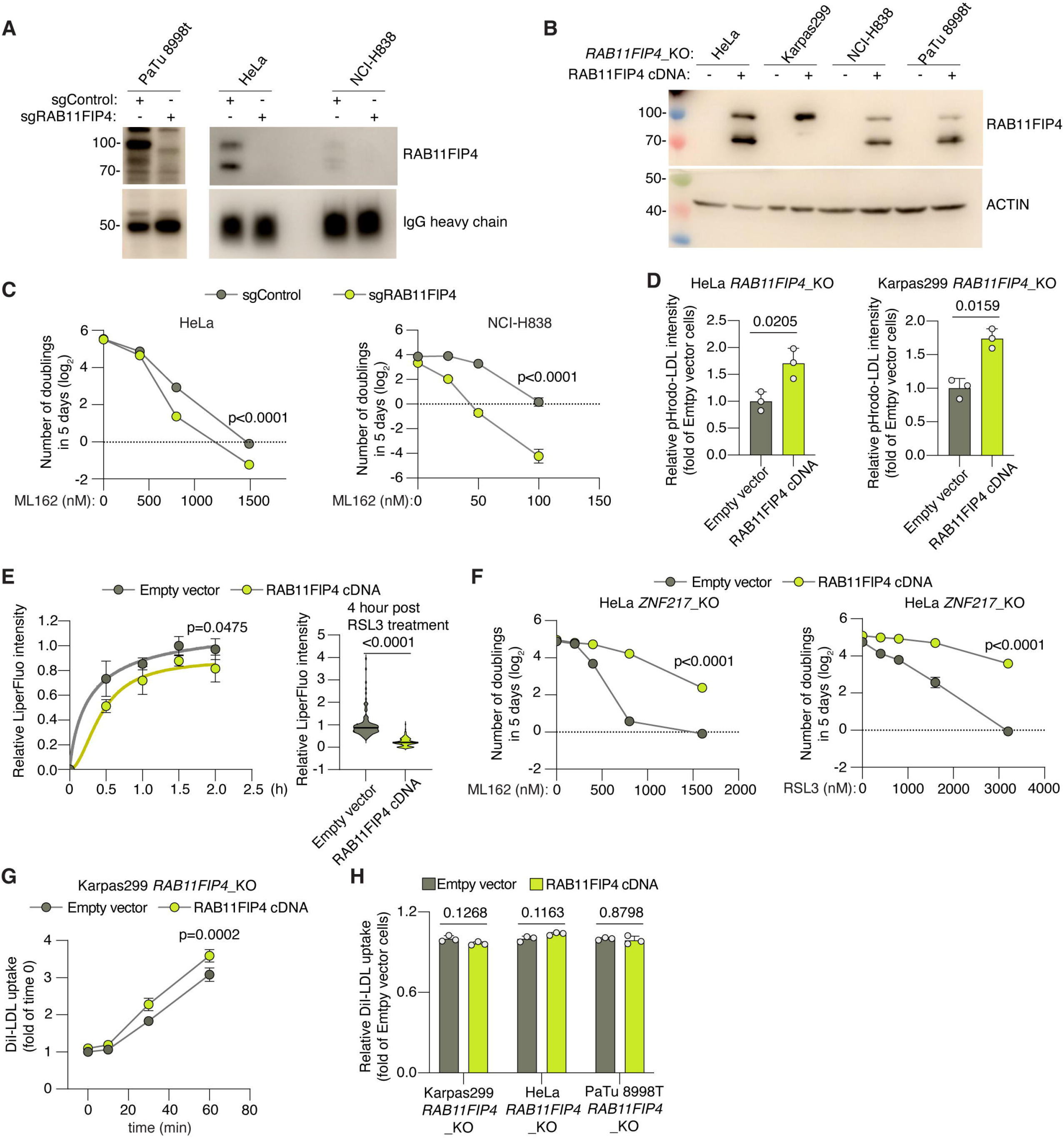
RAB11FIP4 loss sensitizes cancer cells to ferroptosis and impairs lipoprotein assimilation without affecting their uptake. **A,** Immunoblot analysis of RAB11FIP4 in PaTu 8988t, HeLa, and NCI-H838 cells expressing the indicated sgRNAs and after pull down of endogenous RAB11FIP4. **B,** Immunoblot analysis of RAB11FIP4 expression in the indicated *RAB11FIP4*_KO cell lines expressing an empty vector or RAB11FIP4 cDNA. ACTIN serves as loading control. **C,** Number of doublings (log_2_) in 5 days of HeLa (left) and NCI-H838 (right) cells transduced with sgControl or sgRAB11FIP4 after treatment with the indicated concentrations ML162. **D,** Flow cytometry analysis of pHrodo-LDL fluorescence in HeLa (left) and Karpas299 (right) *RAB11FIP4*_KO cells expressing an empty vector or RAB11FIP4 cDNA. **E,** Quantification of lipid oxidation by LiperFluo fluorescence in HeLa *RAB11FIP4*_KO cells expressing empty vector or RAB11FIP4 cDNA at the indicated times after RSL3 treatment (2 μM). **F,** Number of doublings (log_2_) in 5 days of HeLa *ZNF217*_KO cells expressing an empty vector or RAB11FIP4 after treatment with the indicated concentrations of ML162 (left) or RSL3 (right). **G,** Quantification of DiI-LDL uptake at the indicated time points by live-cell imaging in Karpas299 *RAB11FIP4*_KO cells expressing an empty vector or RAB11FIP4 cDNA. **H,** Flow cytometry analysis of DiI-LDL uptake in three *RAB11FIP4*_KO cell lines expressing an empty vector or RAB11FIP4 cDNA. Data in C–H are mean ± s.d.; n = 3 biologically independent experiments. Panels A and B are representative immunoblots. Statistics were performed using two-sided unpaired t tests unless otherwise indicated. Data in C, E, F, and G were analyzed by two-way ANOVA followed by Šídák’s multiple-comparison test.

**Supplementary Figure 5.**
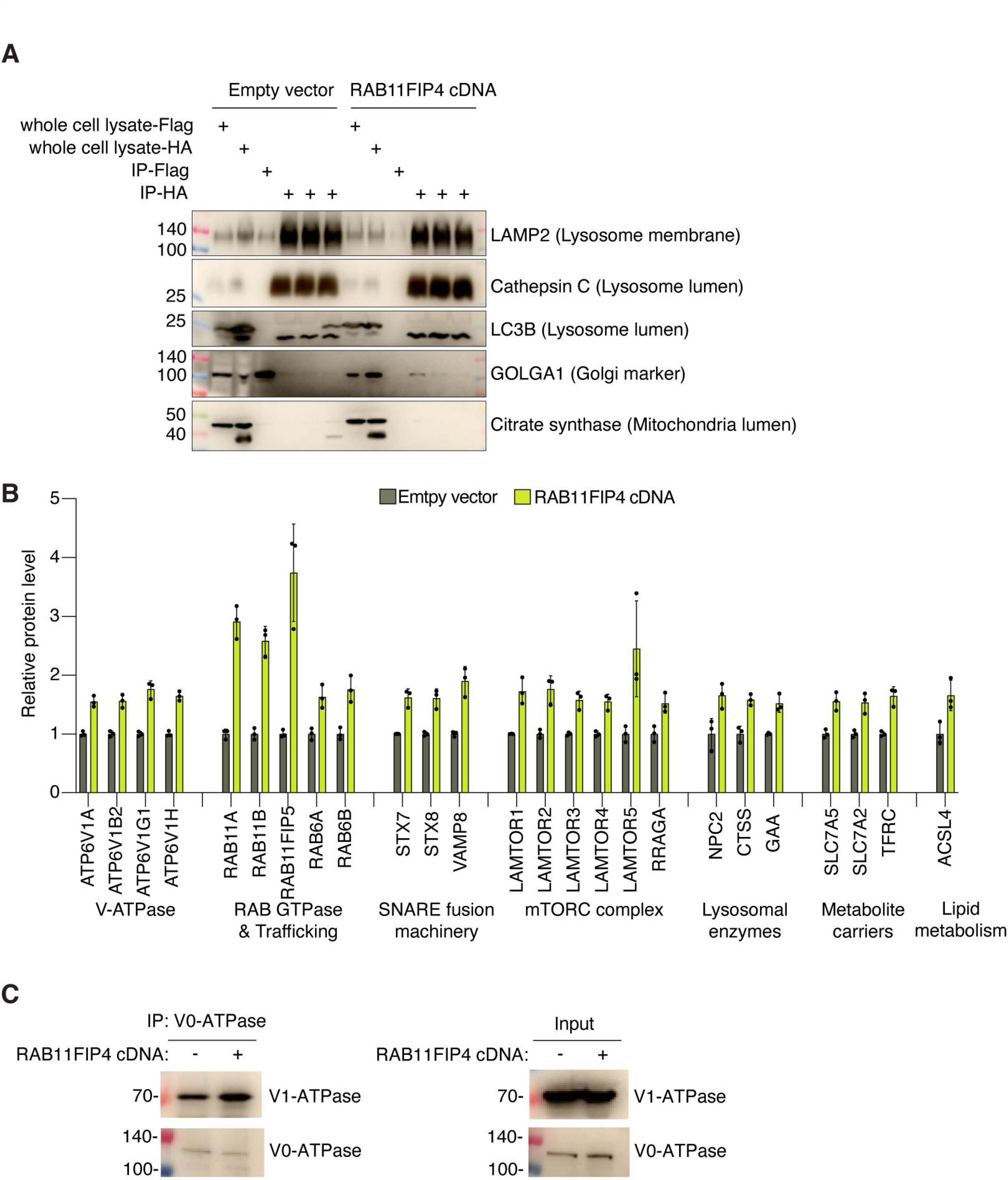
Organellar proteomics identifies RAB11FIP4-dependent changes in the lysosomal proteome. **A,** Immunoblot analysis of LAMP2, cathepsin C, and LC3B showing lysosomal fraction enrichment in whole cell, control IPs (IP-FLAG) and Lyso-IP fractions (IP-HA) from Karpas299 *RAB11FIP4*_KO cells expressing empty vector or RAB11FIP4 cDNA. GOLGA1 and citrate synthase were used as markers of Golgi and mitochondrial fractions, respectively. **B,** Relative levels of indicated proteins identified as differentially expressed in Lyso-IP fractions between Karpas299 *RAB11FIP4*_KO cells expressing empty vector or RAB11FIP4 cDNA. **C,** Immunoblot analysis of immunoprecipitates of the V0 subunit of the lysosomal V-ATPase and its association with the V1 subunit in HeLa *RAB11FIP4*_KO cells expressing empty vector or RAB11FIP4 cDNA. All panels are representative of three independent experiments.

**Supplementary Figure 6.**
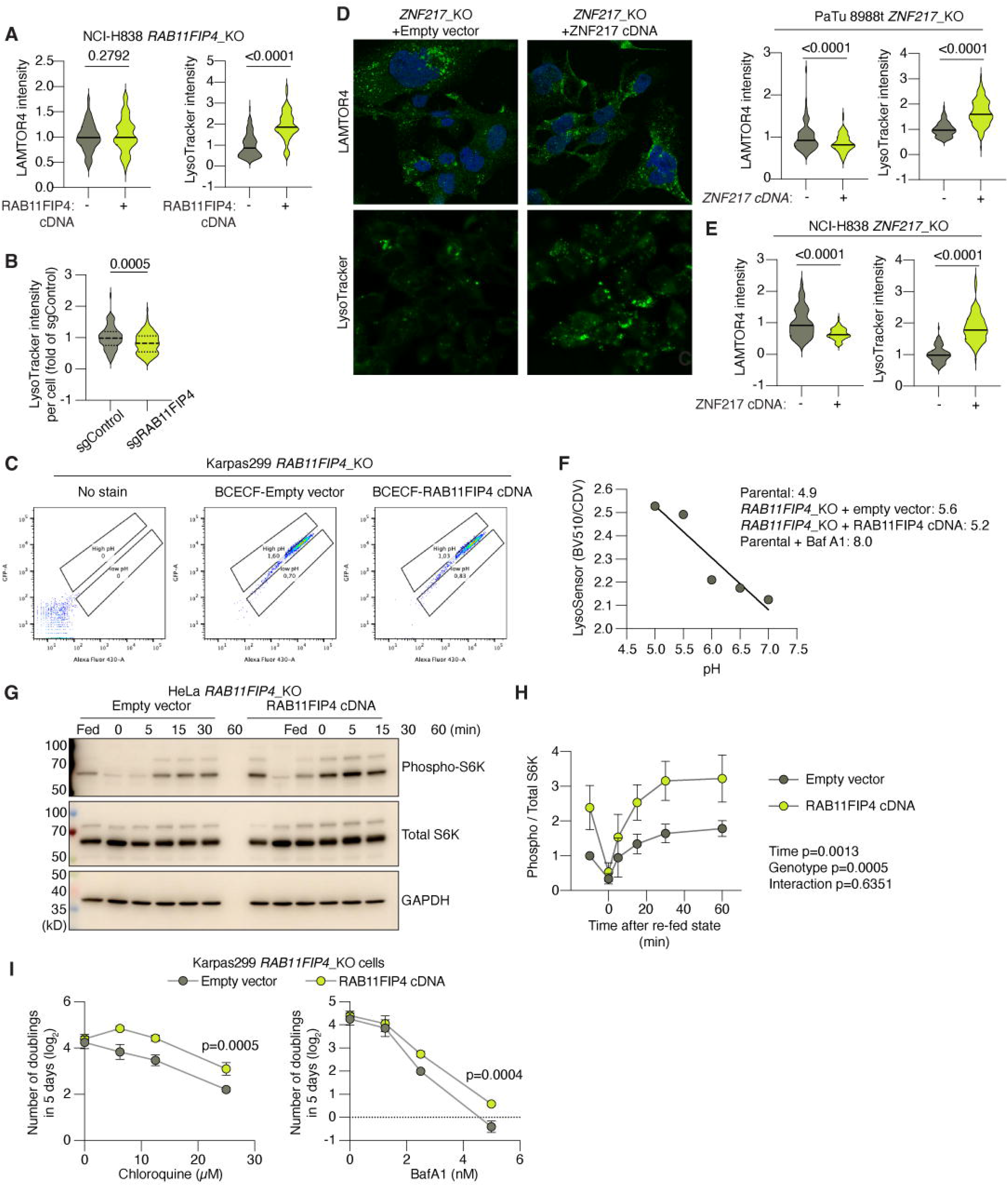
Loss of RAB11FIP4 or ZNF217 mildly decreases cancer lysosomal acidification. **A,** Quantification of LAMTOR4 immunofluorescence (left) and LysoTracker fluorescence in live-cell imaging (right) of PaTu 8988t *RAB11FIP4*_KO cells expressing empty vector or RAB11FIP4 cDNA. **B,** Quantification of LysoTracker fluorescence in HeLa *RAB11FIP4*_KO cells transduced with sgControl or sgRAB11FIP4 using an orthogonal live-cell imaging confocal method. **C,** Cytosolic pH in Karpas299 *RAB11FIP4*_KO cells expressing empty vector or RAB11FIP4 cDNA, measured using BCECF and flow cytometry. **D–E,** Representative images and quantification of LAMTOR4 immunofluorescence and LysoTracker fluorescence using live-cell imaging in PaTu 8988t (D) and NCI-H838 (E) *ZNF217*_KO cells expressing empty vector or ZNF217 cDNA. **F,** Quantification of lysosomal pH in Karpas299 parental and *RAB11FIP4*_KO cells expressing empty vector or RAB11FIP4 cDNA using LysoSensor Yellow/Blue DND-160 and compared to a pH standard curve. Calculated pH values for each cell line are indicated, including a condition with pharmacological lysosomal pH inhibition via treatment with bafilomycin A1 (BafA1, 50 nM for 30 min before analysis). **G–H,** Representative immunoblots (G) and quantification (H) of phospho-S6K and total S6K in HeLa *RAB11FIP4*_KO cells expressing empty vector or RAB11FIP4 cDNA during serum starvation and following serum restimulation. GAPDH is the loading control. **I.** Number of doublings (log_2_) in 5 days of Karpas299 *RAB11FIP4*_KO cells expressing empty vector or RAB11FIP4 cDNA following treatment with the indicated concentrations of chloroquine (CQ, left) or bafilomycin A1 (BafA1, right). Data in A–B, D–F, and H are mean ± s.d.; n = 3 biologically independent experiments. Statistics were performed using two-sided unpaired t tests unless otherwise indicated. Data in H and I were analyzed by two-way ANOVA followed by Šídák’s multiple-comparison test.

**Supplementary Figure 7.**
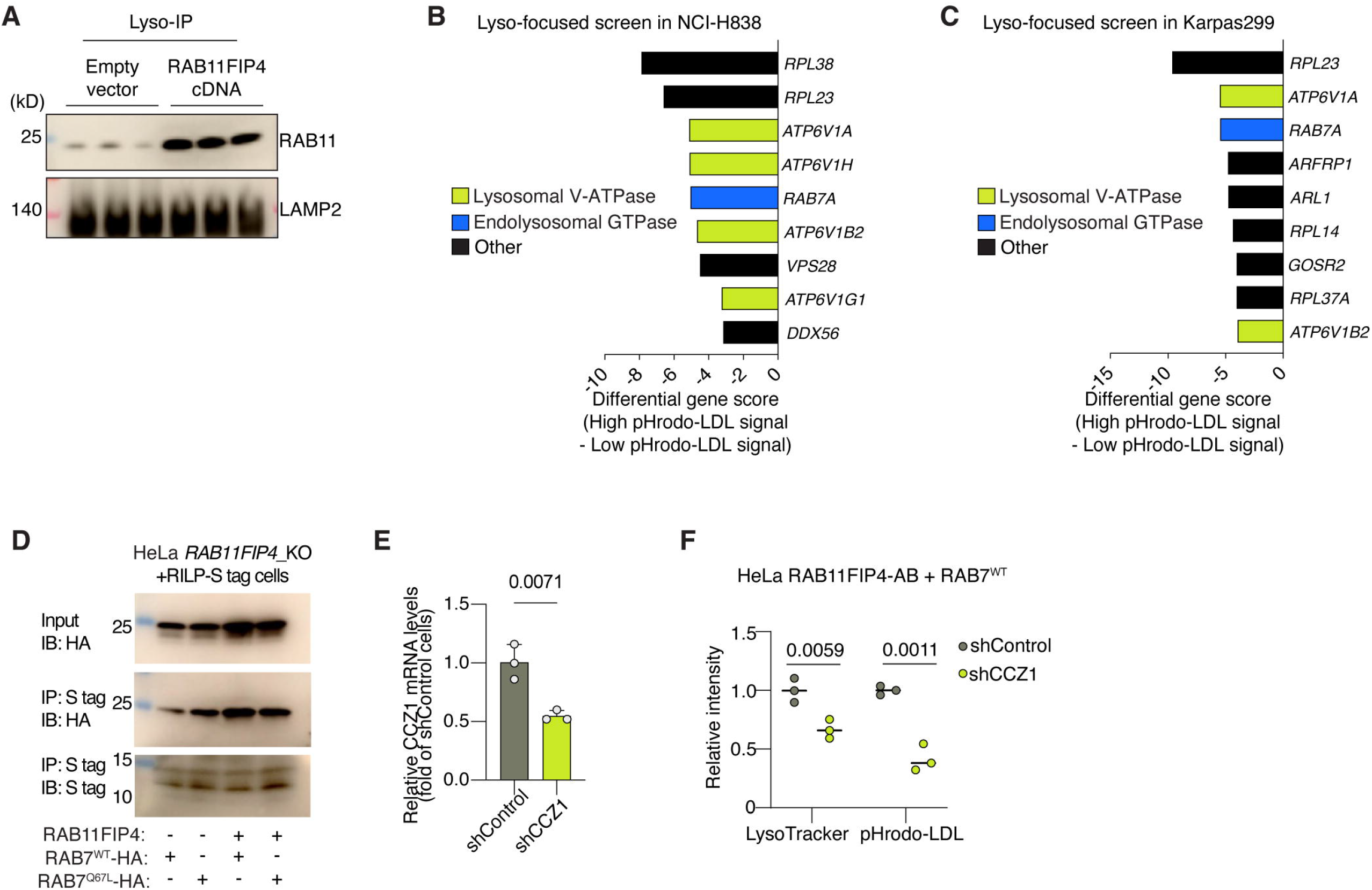
A lysosome-focused genetic screen links the endolysosomal GTPase RAB7 to lysosomal acidification. **A,** Immunoblot analysis of RAB11 in lysosomal fractions (Lyso-IP) isolated from Karpas299 *RAB11FIP4*_KO cells expressing empty vector or RAB11FIP4 cDNA. LAMP2 is used as loading control for lysosomal fractions. **B–C,** Differential gene scores from Lyso-IP-focused pHrodo-LDL CRISPR screens in NCI-H838 (B) and Karpas299 (C) cells. **D,** Immunoblot of RAB7-HA following S-tag pulldown in HeLa *RAB11FIP4*_KO or RAB11FIP4-expressing cells expressing the indicated HA-tagged RAB7 cDNAs and transiently expressing RILP-S-tag. **E,** Relative *CCZ1* mRNA expression in HeLa cells transduced with a control shRNA (shNC) or shCCZ1. **F,** Flow cytometry analysis of LysoTracker and pHrodo-LDL fluorescence in *RAB11FIP4*-expressing HeLa cells transduced with a control shRNA (shNC) or shCCZ1 and expressing RAB7-HA. Data in E and F are mean ± s.d.; n = 3 biologically independent experiments. Statistics were performed using two-sided unpaired t tests unless otherwise indicated.

**Supplementary Figure 8.**
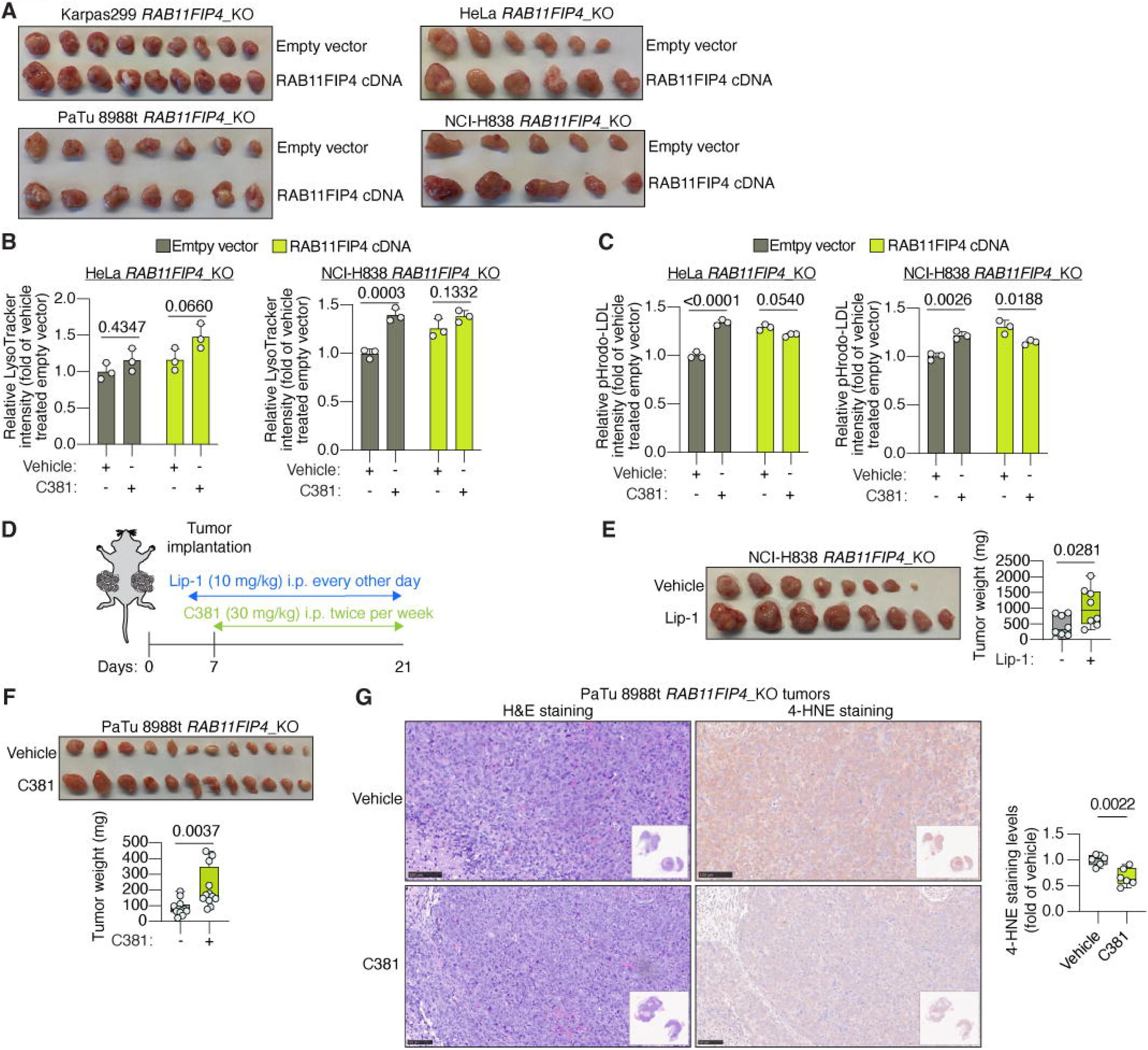
Pharmacological rescue of lysosomal acidification with C381 rescues tumor growth and inhibits tissue lipid peroxidation upon RAB11FIP4 loss. **A,** Representative images of tumors formed by the indicated *RAB11FIP4*_KO cell lines expressing empty vector or RAB11FIP4 cDNA. **B–C,** Flow cytometry analysis of LysoTracker (B) and pHrodo-LDL fluorescence (C) in HeLa (left panels) or NCI-H838 (right panels) *RAB11FIP4*_KO cells expressing empty vector or RAB11FIP4 cDNA after treatment with C381 (10 μM). **D,** Schematic of the experimental approach for Lip-1 and C381 supplementation of tumor-bearing mice. **E,** Representative image (left) and weights (right) of tumors formed by NCI-H838 *RAB11FIP4*_KO cells after supplementation with vehicle or Lip-1 (n = 8 tumors per group). **F,** Representative images (top) and weights (bottom) of tumors formed by PaTu 8988t *RAB11FIP4*_KO cells after supplementation with vehicle or C381 (n = 12 tumors per group). **G,** Representative H&E and 4-HNE immunohistochemistry images (left) and quantification (right) in tumors formed by PaTu 8988t *RAB11FIP4*_KO cells after supplementation with vehicle or C381 (n = 6 tumors per group) In vitro data (B–C) are mean ± s.d.; n = 3 biologically independent experiments. Tumor weights in E–F and 4-HNE quantification in G are shown as box plots; the center line indicates the mean, the box limits indicate the first and third quartiles, and whiskers indicate the range. Statistics were performed using two-sided unpaired t tests unless otherwise indicated. Data in B and C were analyzed by two-way ANOVA followed by Šídák’s multiple-comparison test.

**Supplementary Figure 9.**
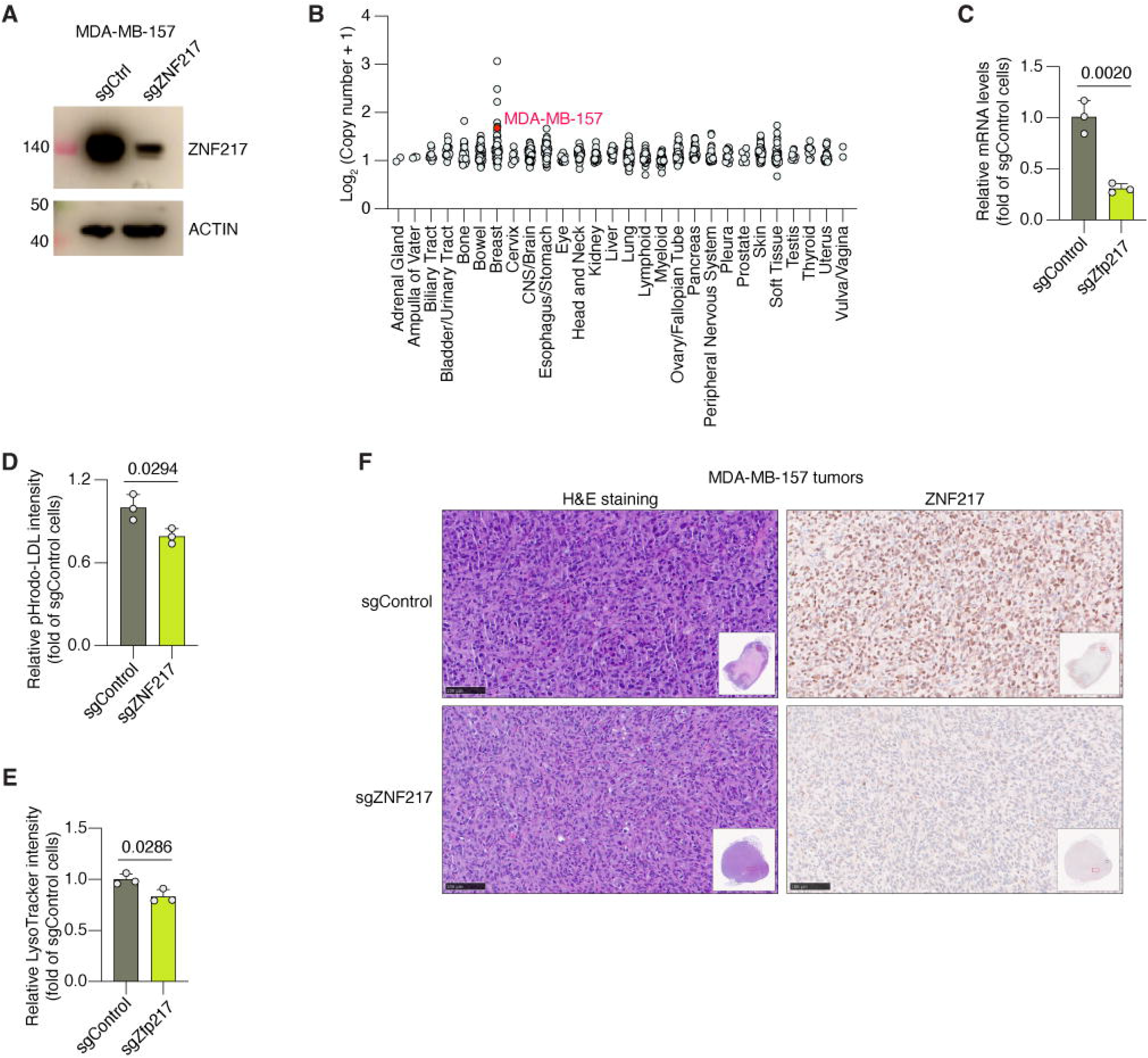
ZNF217 promotes lysosomal function in breast cancer cells. **A,** Immunoblot analysis of ZNF217 in MDA-MB-157 cells transduced with sgControl or sgZNF217. ACTIN is the loading control. **B,** Distribution of *ZNF217* copy number (Log_2_ copy number + 1) across human cancer cell lines from different tissue lineages in the DepMap dataset. Each circle represents an individual cell line. MDA-MB-157 is highlighted in red. **C,** Relative *Zfp217* mRNA expression in murine breast cancer EO771 cells transduced with sgControl or sgZfp217. **D,** Flow cytometry analysis of pHrodo-LDL fluorescence in MDA-MB-157 cells transduced with sgControl or sgZNF217. **E,** Flow cytometry analysis of LysoTracker fluorescence in EO771 cells transduced with sgControl or sgZfp217. **F,** Representative H&E staining and ZNF217 immunohistochemistry of tumors formed by MDA-MB-157 cells transduced with sgControl or sgZNF217. Data in C, D, and F are mean ± s.d.; n = 3 biologically independent experiments. Panels A and E show representative immunoblot and histological images, respectively. Statistics were performed using two-sided unpaired t tests unless otherwise indicated.

**Supplementary Figure 10.**
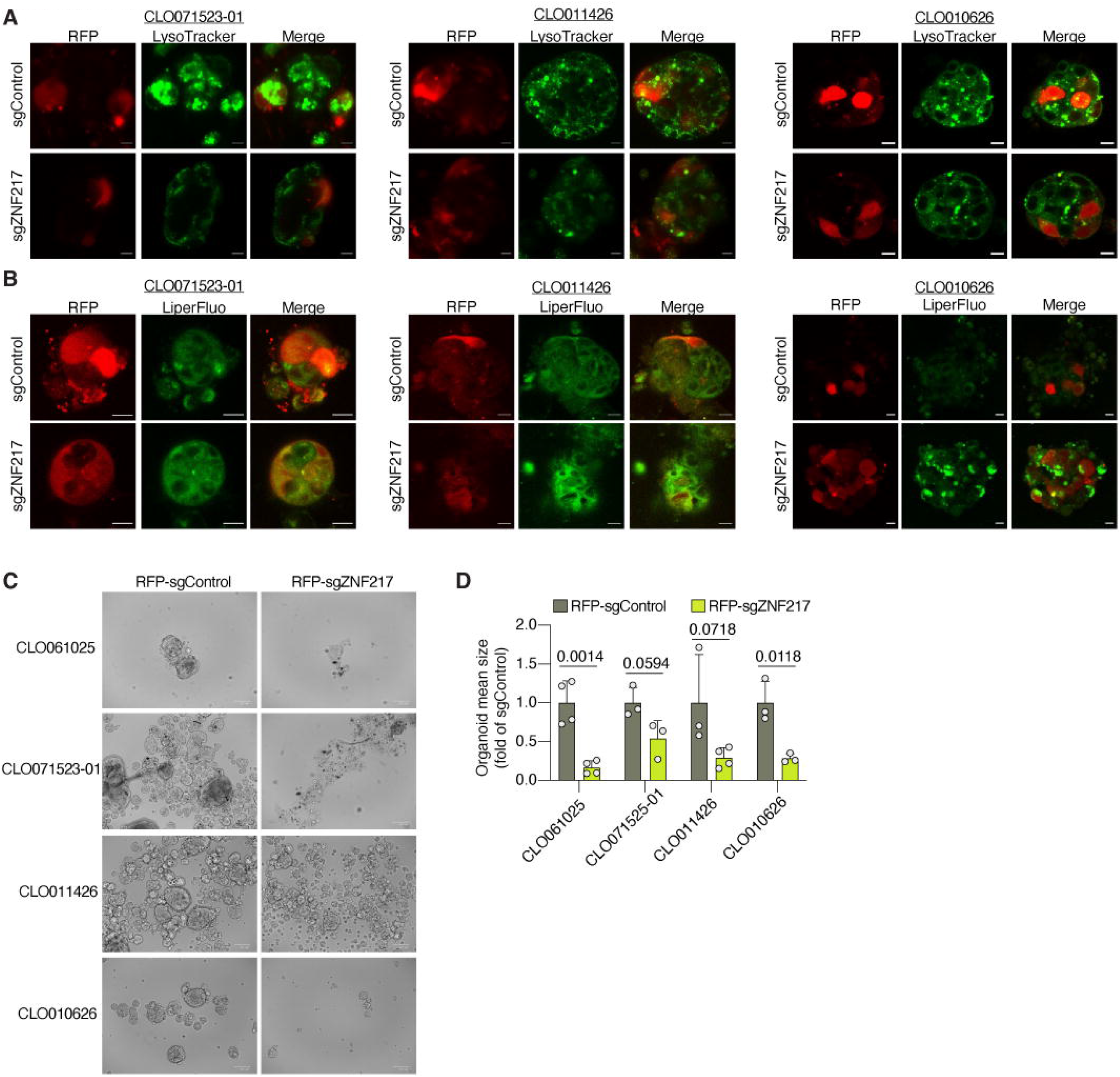
ZNF217 loss reduces lysosomal activity, increases lipid peroxidation, and impairs growth across BC PDOs. **A,** Representative RFP and LysoTracker fluorescence images in indicated PDOs transduced with RFP-sgControl or RFP-sgZNF217. **B,** Representative RFP and LiperFluo fluorescence images in indicated PDOs transduced with RFP-sgControl or RFP-sgZNF217 after treatment with RSL3. **C,** Representative brightfield images of indicated PDOs transduced with RFP-sgControl or RFP-sgZNF217 after 7 days of culture. **D,** Average organoid size of indicated PDOs transduced with RFP-sgControl or RFP-sgZNF217 after 7 days of culture and normalized to size of sgControl-transduced PDO.

**Supplementary Figure 11.**
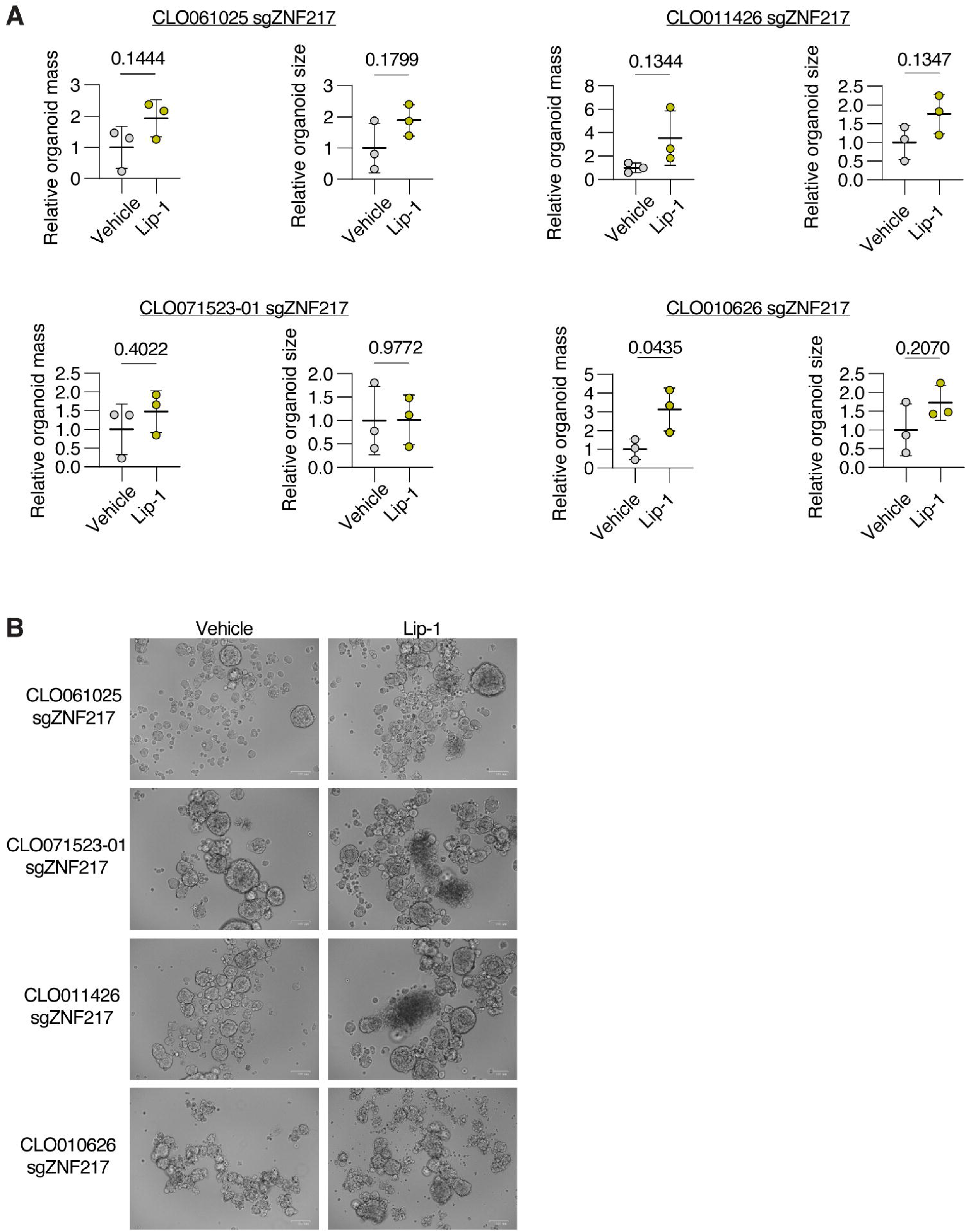
Enhanced ferroptosis partially mediates the growth defects of ZNF217-deficient PDOs. **A,** Quantification of organoid mass and organoid mean size across the indicated sgZNF217-transduced PDOs treated with vehicle or the ferroptosis inhibitor Lip-1 (5 μM) after 7 days of culture. **B,** Representative brightfield images of four sgZNF217-transduced BC PDOs treated with vehicle or the ferroptosis inhibitor Lip-1 (5 μM) after 7 days of culture.

